# Death by a Thousand Cuts – Combining Kinase Inhibitors for Selective Target Inhibition and Rational Polypharmacology

**DOI:** 10.1101/2023.01.13.523972

**Authors:** Ian R. Outhwaite, Sukrit Singh, Benedict-Tilman Berger, Stefan Knapp, John D. Chodera, Markus A. Seeliger

**Author notes:** To whom correspondence should be sent: Markus Seeliger.

## Abstract

Kinase inhibitors are successful therapeutics in the treatment of cancers and autoimmune diseases and are useful tools in biomedical research. The high sequence and structural conservation of the catalytic kinase domain complicates the development of specific kinase inhibitors. As a consequence, most kinase inhibitors also inhibit off-target kinases which complicates the interpretation of phenotypic responses. Additionally, inhibition of off-targets may cause toxicity in patients. Therefore, highly selective kinase inhibition is a major goal in both biomedical research and clinical practice. Currently, efforts to improve selective kinase inhibition are dominated by the development of new kinase inhibitors. Here, we present an alternative solution to this problem by combining inhibitors with divergent off-target activities. We have developed a multicompound-multitarget scoring (MMS) method framework that combines inhibitors to maximize target inhibition and to minimize off-target inhibition. Additionally, this framework enables rational polypharmacology by allowing optimization of inhibitor combinations against multiple selected on-targets and off-targets. Using MMS with previously published chemogenomic kinase inhibitor datasets we determine inhibitor combinations that achieve potent activity against a target kinase and that are more selective than the most selective single inhibitor against that target. We validate the calculated effect and selectivity of a combination of inhibitors using the *in cellulo* NanoBRET assay. The MMS framework is generalizable to other pharmacological targets where compound specificity is a challenge and diverse compound libraries are available.

## Introduction

Biochemical action of pharmacologic agents against unintended targets, or off-target effects, represent a major challenge in biomedical research and medical practice. Although technological limitations once hindered studying drug action beyond a select few biological targets [1], certain drug classes have been found to have more off-targets than anticipated [2-6]. In some cases, inhibitor effects may be due to off-target activities [5]. Some compounds published as selective agents have been found to act through nonselective mechanisms [6], raising concerns about the mechanism of insufficiently validated compounds. Insufficient target validation and off-target effects contribute to clinical attrition rates and the 97% failure rate of compound-interaction pairs brought to clinical trials in oncology [5]. Medicines’ off-target effects, and the symptoms a person experiences following the intended application of a drug, can lead to treatment discontinuance or adverse events [7-9]. Certain symptoms a patient experiences may be directly attributable to action at the intended target of a drug; however, in other cases no clear mechanism may be available, and these symptoms may be due to off-target toxicity.

Kinase inhibitors are a model system for studying target deconvolution and corresponding clinical phenotypes. These inhibitors affect a large class of signaling proteins that are well-studied targets in the treatment of cancers and autoimmune diseases [10]. Certain clinically effective kinase inhibitors are associated with major toxicities which may be due to their off-target effects [11-14]. Recent work suggests that the mechanism of some inhibitors may be due to action against multiple targets or against undefined off-targets even though they may not have been designed as multikinase inhibitors [5, 15].

Clinical efficacy and reduced toxicity could be achieved by rational polypharmacology; simultaneous inhibition of multiple targets after first identifying the most critical targets of a set of compounds. In certain cases, selective inhibition of a single target, as opposed to inhibition of several specific targets, may also be clinically desirable. For example, while Janus kinase (JAK) inhibitors are efficacious in the treatment of rheumatoid arthritis, ulcerative colitis, and other inflammatory conditions [16], existing data highlights how reported rates of adverse medication events and safety profiles differ between JAK-subtype selective medications [14, 17]. *Ex vivo* pharmacokinetics of JAK-family selectivity derived from clinical populations suggests that selectivity for JAK-subtypes may differentiate the safety profiles of JAK inhibitors [18].

Target deconvolution efforts highlight the challenges of navigating the off-target effects of kinase inhibitors [3, 19-27]. Datasets produced by these studies facilitate inhibitor scoring and sorting by chemical selectivity for single kinase targets, and classification of kinases by inhibitor phenotypes [19-21, 23, 28-32]. However, these kinome-wide analyses revealed that most inhibitors have at least several major off-targets. For example, the set of 645 inhibitors evaluated for inclusion in the Published Kinase Inhibitor Set 2 [24], herein referred to as PKIS2, contains many selective kinase inhibitors. These inhibitors were screened at 1 µM in a competitive displacement assay, and the subsequent percent displacement was reported as the action (PKIS2-activity) of the inhibitors. PKIS2 inhibitors exhibit relatively high effects at more than one kinase target compared to the target they show the highest PKIS2-activity against, although interpretation of the relative potency of potent inhibitors using PKIS2-activity values is hampered by the 100% cutoff at the singlicate screening concentration of 1 µM (Fig 1b).

**Fig 1.**
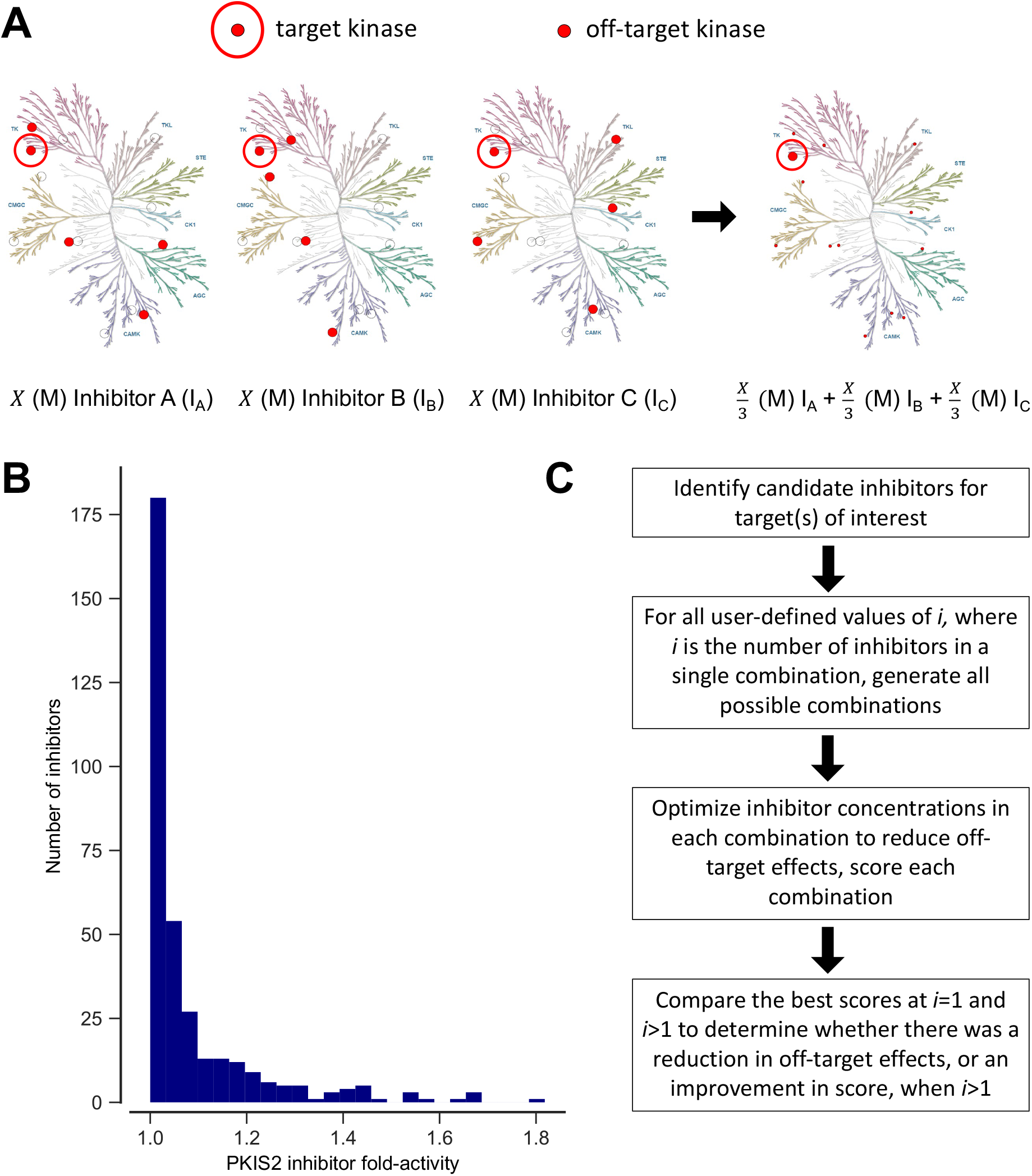
The MMS framework aims to minimize off-target effects. (A) Illustrative K_i_ profiles of three simulated inhibitors against human kinases with shared and equal K_i_s against a target of interest but different off-targets; a combination of the three inhibitors with a cumulative effective K_i_ equal to that of each of the three individual inhibitors retains on-target effect while reducing all off-target effects. In this example, since the inhibitors have equal K_i_s against the target kinase and non-overlapping off-target profiles with equal effects at each off-target, the combination would contain an equimolar mixture of the three single inhibitors. Illustration reproduced courtesy of Cell Signaling Technology, Inc. (www.cellsignal.com) [36]. (B) Fold-activity of inhibitors considered for inclusion in PKIS2. Fold-activity is the PKIS2-activity of each inhibitor against the kinase target it is maximally active against versus the target it is second most active against when screened at 1µM in the DiscoverX KINOMEscan competitive displacement assay. Inhibitors that target multiple kinases with 100% PKIS2-activity are excluded from the analysis. (C) A generalizable framework selects the optimal combination of inhibitors that reduce off-target effects while inhibiting a target of interest.

Several methods measure the selectivity of single compounds, and operational scoring of selectivity varies between methods [20, 23, 28, 30, 31, 33, 34]. The partition index [30], CATDS score [23], or entropy score [31] assigned to an inhibitor are relative selectivity terms describing inhibitor behavior at a target versus a set of off-targets, and the target-specific selectivity score extended this idea for explicit kinase targets [35]. While relative selectivity measures are useful tools, scoring is optimized through both minimization of off-target and amplification of on-target effects. Changes in the off-target distribution could skew towards higher off-target effects in a subset of cases without explicit minimization of off-target behavior. A compound may not be useful if it has a small set of high off-target effects, even if it is relatively selective given the full range of possible off-targets. Target-specific selectivity scoring attempts to address this issue by combining local and global measures of potency that can be controlled with user-defined weights, but ultimately is still a measure of relative selectivity [35]. Conversely, Gini coefficients [28] do not require a defined target, and describe inhibitor behavior against all members of a set. However, Gini coefficients face a similar limitation: they do not discriminate between off-target assignments that build distributions with equivalent areas. A window score [34] or S-score [20] integrate user-defined compound affinity ranges, since only certain magnitudes of off-target effects may be meaningful in context. However, these scoring methods cannot differentiate between the lowest and highest off-target effects within thresholds.

Our goal is to score selectivity for mixtures of inhibitors, unlike current single-inhibitor scoring methods. We consider the design of a selectivity measure that could be applied to both single inhibitors and combinations of inhibitors, and would build upon the strengths of prior measures. This method would define a target or set of targets like relative selectivity measures. It would also weight different ranges of off-target behaviors. However, unlike relative selectivity measures, this ideal measure would maintain sufficient on-target activity without affecting its scoring function in order to optimize inhibitor selection to reduce off-target effects. To our knowledge, no currently available method can define the most selective combination of inhibitors against a single target, or the most selective combination of inhibitors against multiple targets.

Here, we present the multicompound-multitarget scoring (MMS) method for reducing off-target effects by optimizing for inhibitor selectivity using chemogenomic data of kinase inhibitor activity. Inhibitors that act against a common target can be used in fractional combinations, preserving activity at target(s) of interest while reducing off-target effects (Fig 1a; Fig 1c). Our scoring function is only sensitive to changes in off-target effects, weights across a range of off-target effects, and maintains on-target activity without affecting scoring. Integrating current and future chemogenomic datasets will allow experimentalists to predict combinations of inhibitors to more selectively inhibit single targets, or determine the most selective combination of inhibitors for co-inhibition of multiple targets. Determining the most selective combination of compounds with which to co-inhibit multiple targets would represent a significant advance in the implementation of rational polypharmacology towards target inhibition. The translational utility of this method will grow as the set of kinase inhibitors approved for clinical use increases. The MMS method is applicable to any drug-target interaction space and may inform future strategies to reduce off-target effects.

## Methods

### Design of a multicompound-multitarget scoring method to improve the selectivity of target inhibition using inhibitor combinations

Our multicompound-multitarget scoring (MMS) method to calculate the most selective combination of inhibitors against a target set works on multiple data types, is adjustable for certain thresholds of target and off-target inhibition, and has a uniquely flexible scoring framework to meet user-specific goals (Fig 2). MMS specifically optimizes scoring for the reduction of off-target effects rather than a relative change versus the effect at the on-target set. The resulting value known as the Jensen-Shannon Distance (JSD) score is a new metric for inhibitor selectivity that can also describe the total selectivity of a combination of inhibitors against a target set.

**Fig 2.**
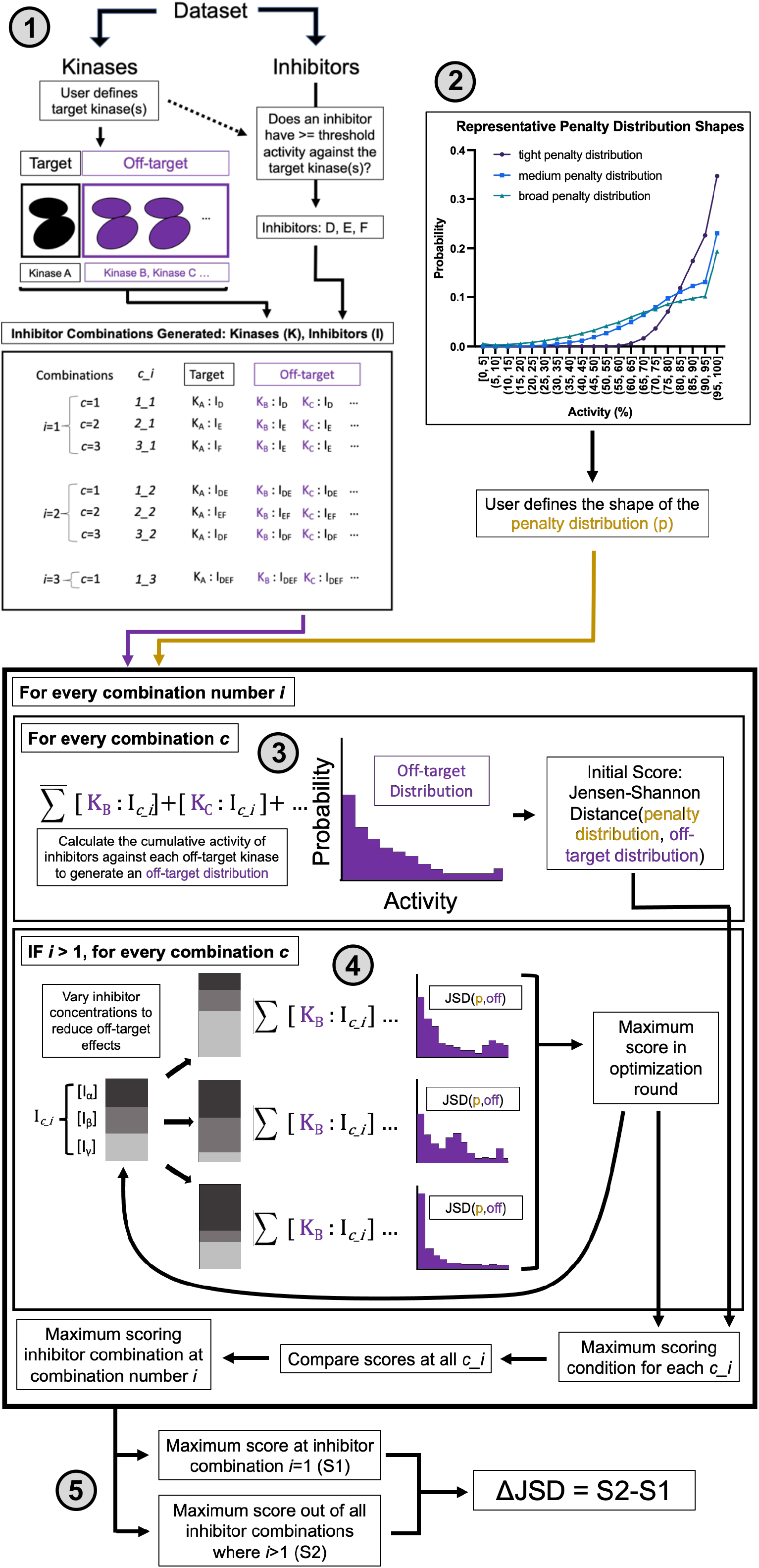
Minimizing off-target effects through optimization of off-target inhibitor activity distributions. (1) All possible inhibitor combinations are generated that maintain activity against a target kinase of interest, up to a maximum user defined value of *i* (the number of inhibitors in a single combination), and for all *c* (a single unique combination for each *i*). Inhibitors that exceed a user-defined minimum activity threshold, defined as ≥ 90% for all analyses presented here, are diluted down to an effective calculated concentration that produces 90% on-target activity. (2) The user defines a penalty prior, a left-skewed probability distribution that is sampled from to build a penalty distribution, which downweighs higher off-target effects more than lower off-target effects. Restricting analyses to one of three shapes; a tight, medium, or broad penalty prior, maintains consistency across experiments. (3) The cumulative activity of inhibitors in a single combination (c*_i*) is calculated for each off-target kinase. A normal distribution is centered on the calculated value to account for possible error in original data measurements. Increasing the variance of this distribution approximates more uncertainty around the original measurements. Distributions from all off-target kinases are used to bootstrap noise for all off-target kinases, which is summed to generate the total off-target distribution, reflecting the activity of the inhibitor combination against all off-target kinases. The Jensen-Shannon distance (JSD score) between the off-target probability distribution and the penalty distribution is calculated: a single metric that describes the distribution of off-target effects. Off-target distributions are scored higher when they are less similar to the penalty distribution, or have fewer off-target effects in the ranges most penalized. (4) The concentrations of inhibitors in a single combination (*c_i*) are optimized to produce the greatest JSD score. (5) The best-scoring combination at every value of *i* is retrieved for all *i*>1, and compared to the *i*=1 condition. If JSD_(*i*>1)_ > JSD_(*i*=1)_, defined by both a statistical increase across technical replicates as well as an absolute threshold, then that combination of inhibitors, at their defined concentrations, is considered to reduce off-target effects relative to the most selective single inhibitor.

If a proposed combination of inhibitors improves the JSD score over a single inhibitor, then that combination will reduce off-target effects relative to that single inhibitor. We use the term activity to describe a measure of target engagement that is nonlinear with respect to K_d_, K_d_^app^, K_i_, or EC_50_ values, the interpretation of which depends upon the input data supplied to the method. For example, K_d_, K_d_^app^, K_i_, or EC_50_ values are used to calculate what the effect of a compound would be if used at an indicated concentration, while PKIS2-activity values are first used to estimate K_d_s for the purpose of calculating the cumulative activity of a compound combination.

The MMS method is summarized here and detailed in our supplemental materials. First, a user-defined selectivity penalty distribution (penalty prior) is chosen to penalize off-target effects across different values of off-target activity. The shape of this distribution affects the penalty assigned to ranges of off-target effects. We recommend using at least two complementary prior shapes in order to gauge what off-target effects can be corrected. We then compare the Jensen-Shannon distance (JSD score) between the single most selective inhibitor (*i*=1) for a target and the combination of inhibitors identified (*i*>1), at optimized concentrations, for that same target. A positive JSD score represents a greater difference between the off-target distribution and the penalty distribution, denoting an improvement in the selectivity of on-target inhibition. We determine significance using both statistical and absolute-difference cutoffs (see supplementary methods).

The data types currently implemented in our MMS approach are built on certain limitations and assumptions. First, chemogenomic data are often derived from kinobead competition-based assays; inhibitors are assumed to be competitive inhibitors. Second, we make reasonable substitutions of K_d_, K_d_^app^, or NanoBRET EC_50_ values for K_i_ values in cumulative activity calculations when K_i_ values are not available. However, we observe that other data types that do not describe target engagement, such as IC_50_ values from cellular proliferation assays, would not be appropriate substitutions for calculating cumulative compound activity. Additional assumptions regarding the effect of target inhibition on the observed phenotype would need to be modeled. Third, kinobead assays may not fully capture kinase dynamics *in vivo*. For this reason, calculations using these data should not be interpreted as a definitive prediction of *in vivo* kinase inhibition when treated with inhibitors. Fourth, kinetic effects of target engagement, such as target-specific phenotypes due to slow off-rates, are not considered. Lastly, unlike datasets that contain K_i_, K_d_, K_d_^app^, or EC_50_ values, performing inhibitor concentration optimization based upon singlicate screen PKIS2-activity values may have more false positives and negatives, and the measurements may not be as precise. A difference of a few PKIS2-activity-scale percentage points for inhibitors in the 90-100% range would produce fold-change differences in calculated dilutions once the inhibitors are recalibrated to minimal 90% target activity. Consequently, we use equimolar ratios of inhibitors in combinations and do not perform these concentration optimization steps in our analyses with PKIS2 inhibitors based on singlicate screen data. For example, given a reference concentration of 1 µM, a combination of two inhibitors would be used at 500 nM each. We also include an additional high off-target penalty parameter that can be adjusted (supplementary methods).

JSD scoring during a set of technical replicates of the MMS method with the same input parameters is limited by the scope of a single penalty prior shape. Scoring can be tuned to the dynamics of both the system in question and the goals of the user by altering the shape of the penalty prior. For this reason, we use a standardized pair of complementary penalty priors across all analyses: the tight penalty prior with either the medium or broad penalty prior (Fig 2). It is possible that different inhibitors are identified as the most selective single inhibitor, or combination of inhibitors, for the same target(s) when using different penalty priors, although the initial inhibitor or inhibitor set may also lead to significant reduction in off-target effects.

The MMS method works on datasets containing K_i_, K_d_, K_d_^app^, EC_50_ or PKIS2-activity values. Additional datatypes could be integrated into future extensions of this approach with appropriately modeled compound-target interactions. The penalty prior is a flexible tool to meet user-specific goals, and differentially weighs off-target effect ranges unlike other selectivity metrics. The JSD scoring metric integrated in our method is only sensitive to changes in the off-target distribution, and our inhibitor concentration optimization protocol minimizes off-target effects while retaining functionally meaningful on-target activity. MMS calculates the most selective single inhibitor against a single target kinase, yielding a new selectivity metric for describing single inhibitors. In addition, MMS can also calculate the most selective combination of inhibitors against a single target kinase, or even multiple target kinases.

## Results

### Inhibitor combinations outscore single inhibitors with increasing inhibitor set size

Kinase-inhibitor data for sets of 100 kinases are simulated from the selectivity profiles of the average inhibitor in PKIS2, less selective inhibitor sets with equivalent probabilities of available potent targets above a 90% activity threshold, or binary inhibitors with either potent or zero activity (Fig 3a). Specifically, probability distributions reflecting these four inhibitor types (Fig 3a) were used to sample 100 points for each inhibitor that was simulated in each artificial dataset, reflecting the activity of that inhibitor against each of the 100 kinase targets. Calculation of the dilution of the most selective single inhibitor that maintains 90% target activity, for all 100 simulated kinases, decreases other off-target effects relative to the initial selectivity profile (Fig 3d, Fig S4;S5). Increasing the number of simulated inhibitors increases the selectivity of the most selective single inhibitor, on average, for all targets (Fig S9a). Correspondingly, the average magnitude of improvement was less for targets with highly selective single inhibitors, since JSD scoring is bounded at 1 (Fig S9b).

**Fig 3.**
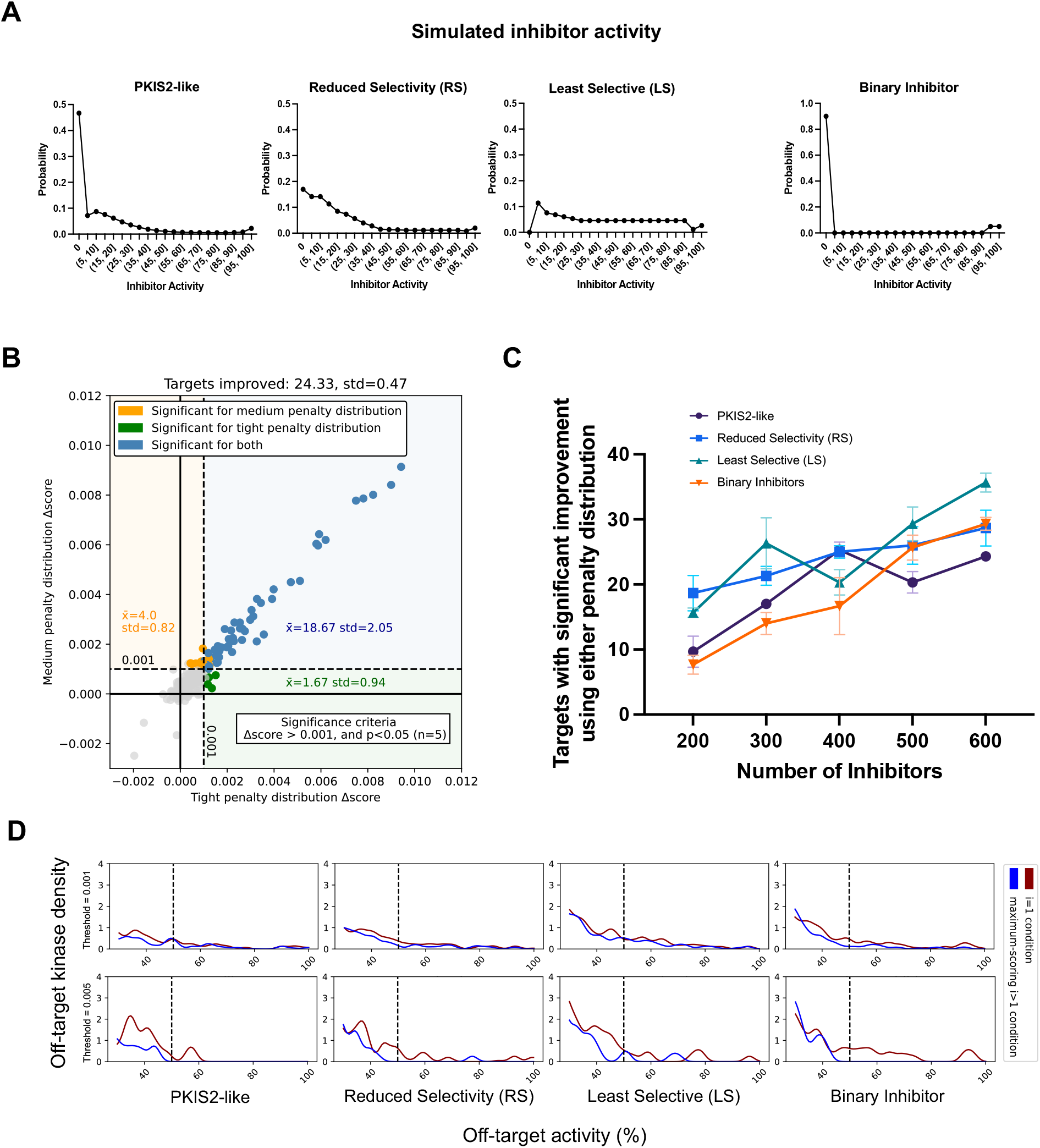
Simulated inhibitor combinations reduce off-target effects for single targets. (A) Simulated activity profiles for four inhibitor types. PKIS2-like represents the average selectivity profile of the 645 compounds screened for inclusion in PKIS2 [24]. This selectivity profile was altered to produce a less selective inhibitor profile (Reduced Selectivity RS), an even less selective inhibitor profile (Least Selective LS), and an inhibitor selectivity profile with either potent or zero activity (Binary Inhibitor). These four activity profiles are used to generate simulated datasets of 100 targets and 200, 300, 400, 500, or 600 inhibitors. The minimum on-target potency required to generate combinations is 90% across all analyses; the probability of ≥ 90% activity for a single inhibitor is approximately equivalent for all PKIS2-like, Reduced Selectivity, and Least Selective inhibitor types, given random sampling from the underlying distributions, ensuring that the probability of generating a possible combination is similar between inhibitor types. The cumulative probability for the binary inhibitor at ≥ 90% activity is 10%. (B) Three independent datasets of 600 PKIS2-like inhibitors against 100 targets are simulated. The number of targets in each dataset with a significant change in score, for the highest-scoring combination of two or three inhibitors versus the most selective single inhibitor, are determined. Significance is defined by both a statistically significant improvement in mean score from the highest-scoring *i*=1 to *i*>1 combination, and an absolute difference in score of at least 0.001, for either penalty distribution type. The average number of targets with a significant improvement for either or both penalty distribution types across the three independent simulations, as well as the average number of targets with any of the three types of significant improvement, are indicated in the figure. (C) Increasing sets of simulated inhibitors are analyzed with MMS. The number of targets that can be more selectively inhibited with greater than one inhibitor relative to the most selective single inhibitor, for either or both penalty distribution types, are indicated. Three independent sets of inhibitors are simulated and analyzed for each type of inhibitor at every dataset size (n=3), with five technical replicates of the MMS analysis performed for each inhibitor set to asses significance. Data represented as mean ± SEM. (D) Density functions illustrate the average off-target effects for the highest scoring single inhibitor (blue) versus the highest scoring combination of inhibitors for targets with significant improvements (maroon), scored using a tight penalty distribution. Vertical dotted lines represent the approximate activity value where the tight probability distribution becomes negligible. The variance of the normal distributions in the figure is the same as scoring method: 2.5, such that the overall density mimics each respective off-target penalty distribution. The subset of targets with an improvement ≥ 0.005 is also indicated in the second row of (D).

Combinations of either two or three PKIS2-like inhibitors (max *i*=3) scored with either the tight or medium penalty priors improve the off-target profiles for 10% of kinases when highly selective PKIS2-like inhibitors are used at an inhibitor:kinase ratio of 2:1 (Fig 3c, Fig S3). Increasing the inhibitor to kinase ratio improves the likelihood of decreasing off-target effects (Fig 3c, Fig S3), despite also increasing the probability of simulating a more selective single inhibitor. The off-target profiles of almost 25% of targets could be improved using an inhibitor to kinase ratio of 6:1 (Fig 3b). Decreasing the selectivity of the simulated inhibitors used also appears to increase the chances of identifying a more selective inhibitor combination than the most selective single inhibitor (Fig S3). Increasing the ΔJSD cutoff to 0.005 enhances the observed effect size for combinations that meet this threshold (Fig S6; S7; S8). These simulated inhibitors are uncorrelated; unlike real inhibitors, there is not an increased likelihood of activity against structurally similar targets. While the likelihood of inhibiting a given target more selectively with a combination is related to combinatorial availability for that target and the off-target space covered by the underlying dataset (Fig S3), the subset of targets that scored most highly with either penalty prior also scored positively and significantly with the other complementary penalty prior shape. Improvement in off-target distributions for simulated targets was dominated by positive ΔJSD scores for both penalty priors used (Fig S3; S6). This suggests that, with ample off-target dataset coverage and inhibitor combinatorial space, the property of observing an overall improvement in off-target effects, a positive and significant ΔJSD score, is not restricted to a specific penalty prior shape. Unlike the ΔJSD score, single kinase off-target profiles, or the set of observed off-target effects, should be interpreted in context of the penalty prior shape used for scoring; it is not uncommon for the off-target profile to increase relative to the most selective single inhibitor in activity ranges with low or zero penalties (Fig S4; S5; S7; S8). These simulated analyses demonstrate that, with increasing dataset size, greater combinatorial space will increase the utility of the method, despite also increasing the probability of identifying more selective single compounds.

### Inhibitor combinations improve selectivity over the most selective single inhibitors for single kinase targets

We consider the optimal combination of inhibitors for all unique single targets in PKIS2 [24] (max *i*=3), Klaeger *et al* [23] with 243 clinically evaluated inhibitors (max *i*=6), Karaman *et al* [20] with only 38 inhibitors (max *i*=5), or Davis *et al* [21] with 72 inhibitors (max *i*=5) (Fig 4). PKIS2 contains 645 inhibitors at a coverage ratio of 1.6:1, and 24, or, 6% of unique targets, can be significantly improved given both statistical and absolute ΔJSD cutoffs (Fig 4, Fig S10; S11). Targets with greater ΔJSD scores have greater improvements in their off-target profiles (Fig S10; S11). These data indicate that global off-target activity may be reduced for a small but not insignificant group of single kinase targets using merely two or three inhibitors instead of the most selective single inhibitor.

**Fig 4.**
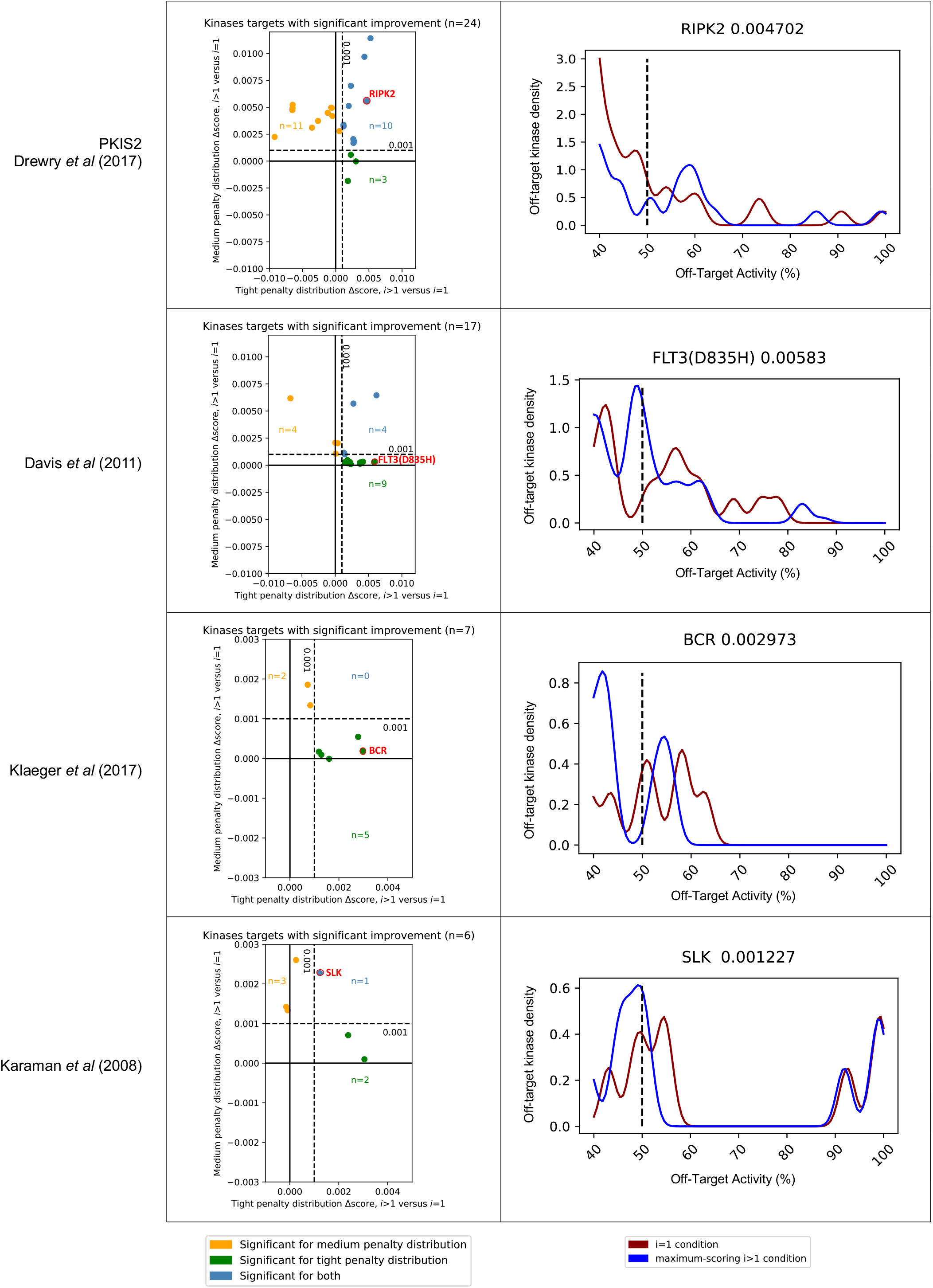
Inhibitor combinations using chemogenomic data reduce off-target effects for single kinase targets. Combinations of two or three inhibitors reduce off-target effects for some kinase targets across three chemogenomic datasets. Scatterplot data is represented as the change in JSD score using a tight penalty prior (x-axis) versus the change in JSD score using a medium penalty prior (y-axis). Five technical replicates are performed for all analyses; significance was determined by an absolute improvement of greater than 0.001 in score, as well as statistical significance between the mean *i*>1 and *i*=1 scores. Probability density functions (pdfs) illustrate the average off-target effects for the highest scoring single inhibitor (blue) versus the highest scoring combination of inhibitors (maroon) for representative examples, scored using a tight penalty distribution. The variance of the normal distributions used to generate the pdfs is the same as was used in the scoring method (2.5) such that the overall density is an accurate reflection of the off-target penalty distributions. Vertical dotted lines represent the approximate activity value where the tight probability distribution becomes negligible.

Limiting the off-target effects against only a subset of kinases might be a user case of interest. The Eph receptor tyrosine kinase family are promising targets in immunotherapy [37]; in the following analysis off-target effects at other kinases are not considered. Some PKIS2 compounds have relatively low PKIS2-activity against EPH kinases, but the flexibility of the MMS method allows such off-target effects to be adequately scored with the broad prior distribution. Four targets, EPHA2, EPHA3, EPHA4, and EPHA5, can all be more selectively inhibited using a combination of inhibitors (Fig S18). This case example illustrates how off-target effects can be studied within kinase families of interest, and how the flexibility of the user-defined penalty priors facilitates specific analyses.

MMS analyses of the Davis *et al* and Karaman *et al* datasets also indicated improvements in selectivity for single targets. A coverage ratio of 0.16:1 across the Davis *et al* dataset yielded improvements for 17 targets (Fig S12; S13), while a coverage ratio of 0.12:1 across the Karaman *et al* dataset, which included only 38 inhibitors, was able to improve 6 targets (Fig S14; S15).

The Davis *et al* dataset contains selectivity screening for kinase mutants in addition to wild-type constructs, including those of ABL, EGFR, FLT3, and KIT. Interestingly, multiple FLT3 alterations were found to be more selectively inhibited using a combination of inhibitors than a single inhibitor, and these inhibitors were different from those which were most selective for FLT3. Although this is a small sample size, it raises the intriguing possibility that this method may be useful in cases where selective single-compound pharmacological targeting of clinically relevant mutations has not yet been actualized.

We investigate whether or not it is possible to reduce the off-target effects of clinically evaluated inhibitors, using Klaeger *et al*, which also contains measurements of inhibitor action against non-kinase off targets. It is possible that the comprehensive methodological determination of K_d_^app^ values, which were derived from measurements of both target engagement and compound method of action in whole cell lysates, may have limited detection of weak compound-target interactions; the data matrix is notably sparse in some areas. PKIS2 assigns a PKIS2-activity value of 0 to 47% of compound-kinase interactions, while Klaeger *et al* does not assign a K_d_^app^ in 94% of cases, limiting the effectiveness of off-target cross-comparison calculations. It is also possible that other methods, such as singlicate screening, over-assign weak compound-target interactions. Nevertheless, single target analysis suggested modest improvements in selectivity were possible for EPHA5, BCR, PDK1, and PRKCD, using a tight penalty prior, all while maintaining on-target activity of at least 90%. While an inhibitor combination resulted in a subtle shift in the off-target profiles of TYK2 and PRKD2, these do not appear to be meaningful changes. Single-target analysis from this dataset suggests that the single most selective inhibitor is generally the most effective.

Differences between assignments from Davis *et al*, Karaman *et al*, and Klaeger *et al* may be due to varied inhibitor set composition, kinase coverage space, or K_d_/K_i_/K_d_^app^ values. No comprehensive cross-analysis of compound activity across such datasets has been performed, and might illustrate variability in observed compound activity due to different experimental techniques.

### MMS prediction of cumulative compound activity is validated in cellulo

We aimed to determine whether the cumulative activity of compounds *in cellulo* would match predicted activity according to our MMS scoring framework. We selected compounds from the PKIS2 dataset in order to evaluate whether singlicate screening PKIS2-activity data could have translational utility. Compounds TPKI-108, UNC10225285A, and UNC10225404A have validated on-target action against MAPK14 (p38-alpha), with experimentally determined K_d_s of 150 nM, 140 nM, and 250 nM respectively [24]. These values matched the MMS 90% activity threshold of ∼111 nM quite well. MMS suggested that TPKI-108 was the most selective compound against MAPK14 (*i*=1), but that a combination of all three could modestly reduce off-target effects.

We studied the activity of these compounds against MAPK14 and major off-targets using the in-cell NanoBRET target engagement assay (Fig 5, Fig S20). We found that the EC_50_s of these compounds diverged from previously observed K_d_s; TPKI-108 was slightly more potent (EC_50_=82 nM), while UNC10225404A (EC_50_=4.6 µM) and UNC10225285A (EC_50_=1.8 µM) were less potent (Fig 5a). Single compound activity at off-targets also varied from singlicate screen PKIS2-activity values. For example, UNC10225404A had high PKIS2-activity against MAPK14 (p38-alpha) >90% and only 18% PKIS2-activity against MAPK11 (p38-beta). Generally, a 30% or 35% threshold is used for further validation of activity screen hits, so an 18% PKIS2-activity measurement would have been excluded from most off-target validation studies using common experimental protocols. NanoBRET data suggested that the potency of UNC10225404A against both proteins was quite similar (MAPK14: EC_50_=4.6 µM, MAPK11: EC_50_=2.4 µM) (Fig 5a). About half (5/11) of the kinase off-targets that were annotated as having high PKIS2-activity at 1 µM with these compounds in the PKIS2 screen were not readily observed off-targets in the NanoBRET assay (EC_50_ > 50 µM for all replicates) (Fig S20). These data suggest that different experimental techniques or the biochemical state of kinase proteins in different experimental formats may produce significant deviations in observed compound-kinase activity measurements, both for singlicate PKIS2-activity data and even multiple-point K_d_ curves. We also conducted NanoBRET target engagement assays using cell lysates in order to validate compound-target interactions past our conservative 50 µM detection cutoff for the *in cellulo* assay (Fig 5b). These data closely matched our *in cellulo* results and provided additional validation of lower potency off-target EC_50_s.

**Figure 5.**
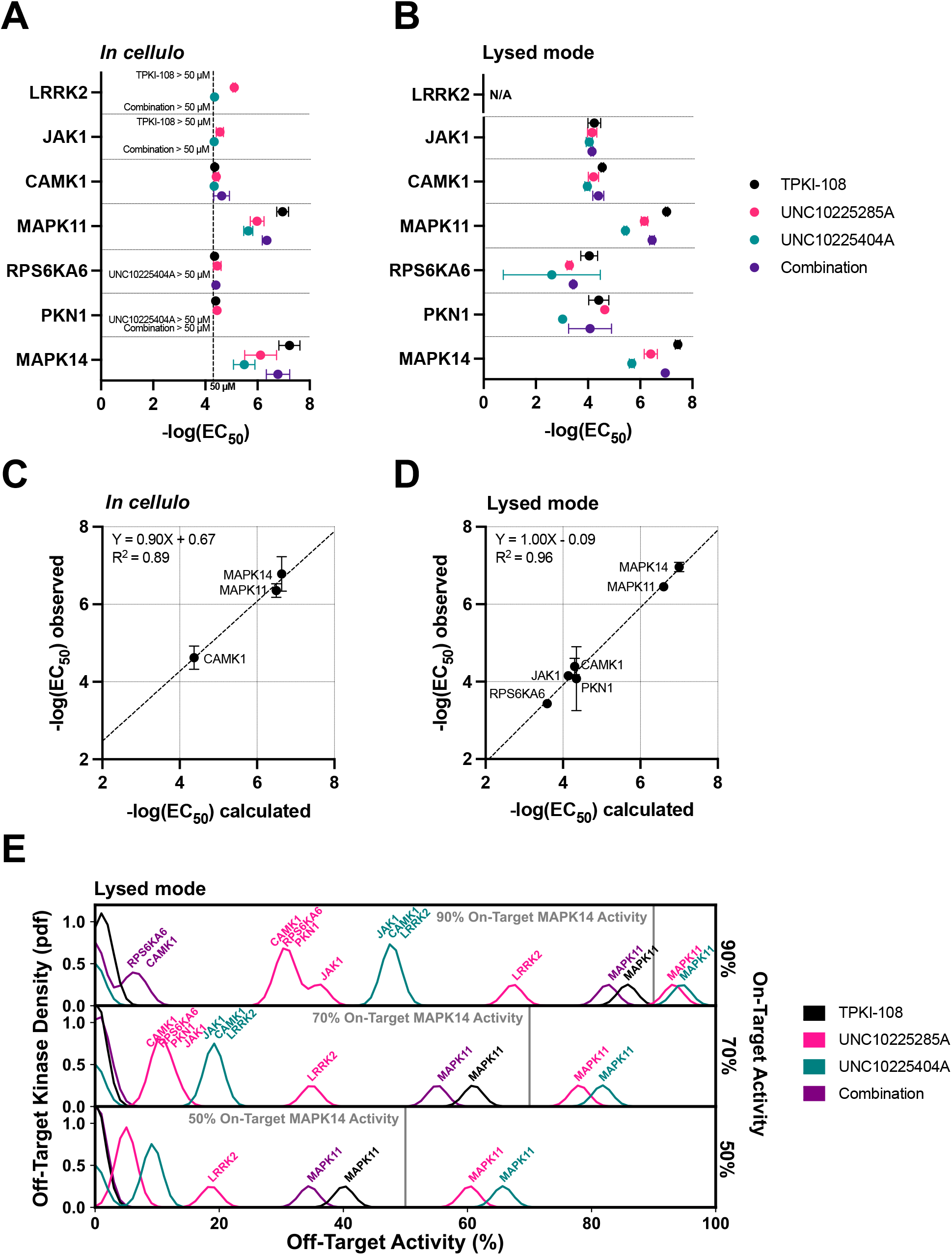
Validation of compound combination predicted activity and selectivity. The cumulative activity of each compound and the equimolar combination of compounds was studied with the 11-point in-cell NanoBRET target engagement assay *in cellulo* (A, C) and using cell lysates (B, D). EC_50_s ± std were obtained for MAPK14 and those off-targets with high PKIS2-activity for any of the three single inhibitors (A, B) Five replicates of the *in cellulo* experiments were performed and two replicates of the lysed mode experiments were performed; only those replicates for each kinase that could be detected at a cutoff of 50 µM were included in calculations (Fig S20). The predicted cumulative activity of the three compounds, in their equimolar mixture, was calculated from the average EC_50_ values of the individual compounds using the same protocol in the MMS method and matched observed EC_50_s (C, D). Simple linear regressions were performed for both *in cellulo* and lysed mode data (C, D). Pearson correlations were significant for the *in cellulo* inhibitor combination EC_50_ calculations and observations (R^2^=0.98, p < 0.05, one tail) and were highly significant (R^2^=0.99, p < 0.0001, one tail) for the lysed mode EC_50_ calculations and observations. Calculation of the off-target profiles of the inhibitor combination and the individual inhibitors suggests that the combination modestly improves selectivity for MAPK14 over the most selective single inhibitor, TPKI-108 (E). The off-target activity of the inhibitors is calculated based upon their EC_50_s (lysed mode) for each off-target kinase, at the concentration needed to reach 90%, 70% or 50% on-target activity against MAPK14. The variance of the normal distributions used to generate probability density functions (pdfs) is the same as was used in the MMS scoring method (2.5) and the activity scale is shown between 0 and 100.

Notably, the cumulative activity of the three compounds dosed in equimolar ratios matched what would have been expected from the single-compound NanoBRET EC_50_ values in both the *in cellulo* and lysed mode assays (Fig 5c; Fig 5d; Fig S21). For example, given the average three *in cellulo* EC_50_ values of the single inhibitors against MAPK14, the cumulative activity of the equimolar combination is calculated to be EC_50_=231 nM, very close to the average observed EC_50_=246 nM. These calculations were performed using the same protocol as in the MMS method (see supplementary methods, Fig S21). This excellent matching was observed for the three kinases with sufficient EC_50_ data for performing calculations from the *in cellulo* experiments, and for all kinases from the lysed mode experiments. These observations strongly support our hypothesis that the cumulative activity of a combination of compounds can be predicted given the activity of each single compound against both targets and off-targets of interest. Since NanoBRET EC_50_s measure target engagement, as opposed to IC_50_s generated from commonly performed cellular proliferation assays that observe cellular phenotypes induced by target inhibition, these are more reliable form of EC_50_ data for calculating cumulative target activity.

We used the EC_50_ values from the *in cellulo* or lysed mode experiments to calculate what the off-target effects of the different inhibitors would be if used at the specific concentrations necessary to produce either 90%, 70%, or 50% on-target activity against MAPK14 (Fig 5e; Fig S22). We find that there is a modest increase in the selectivity of MAPK14 inhibition when using the combination of inhibitors over the most selective single compound, TPKI-108, which is already highly selective. Although minor off-target effects are introduced as a result of adding more inhibitors, as expected, the combination has less activity against the primary off-target, MAPK11, compared to TPKI-108.

These proof-of-concept experiments support our hypothesis that combinations of inhibitors can improve the selectivity of target inhibition by reducing high off-target effects. Even if such combinations add more unique off-targets, by dosing the compounds at the appropriate concentrations we expect that these minor off-target effects will be negligible. We observed that EC_50_ values in the NanoBRET format varied from K_d_ values and PKIS2-activity values determined in a separate experimental system. While the combination still modestly improved selectivity over the most selective single compound, our NanoBRET data (Fig 5c; Fig 5d) suggest that predictions of cumulative compound activity would be most accurate if generated from single-inhibitor data derived from the same experimental system. Our combination of three inhibitors modestly improved selectivity over the already highly selective inhibitor TPKI-108. We expect that, for targets that lack highly selective single inhibitors, or in cases of multiple on-targets, the magnitude of improvement in on-target selectivity will be greater.

### MMS enables rational polypharmacology by nominating inhibitor combinations

To our knowledge, no method has yet been developed to determine the most selective single inhibitor, or set of inhibitors, given the cumulative effects of those drugs at multiple targets and off-targets. Such a method may be of particular use to those who wish to target compensatory pathways or multiple components of the same pathway. Cases of multiple on-target kinases are considered; the most selective combination of inhibitors for multiple targets is not necessarily the combination of the single most selective inhibitors at each target. The kinase pairs YES/FES and ROCK1/ROCK2 are selected as case examples from the PKIS2 dataset (Fig 6, Fig S19). YES and FES are nonreceptor tyrosine kinases with divergent inhibitor phenotypes.

**Fig 6.**
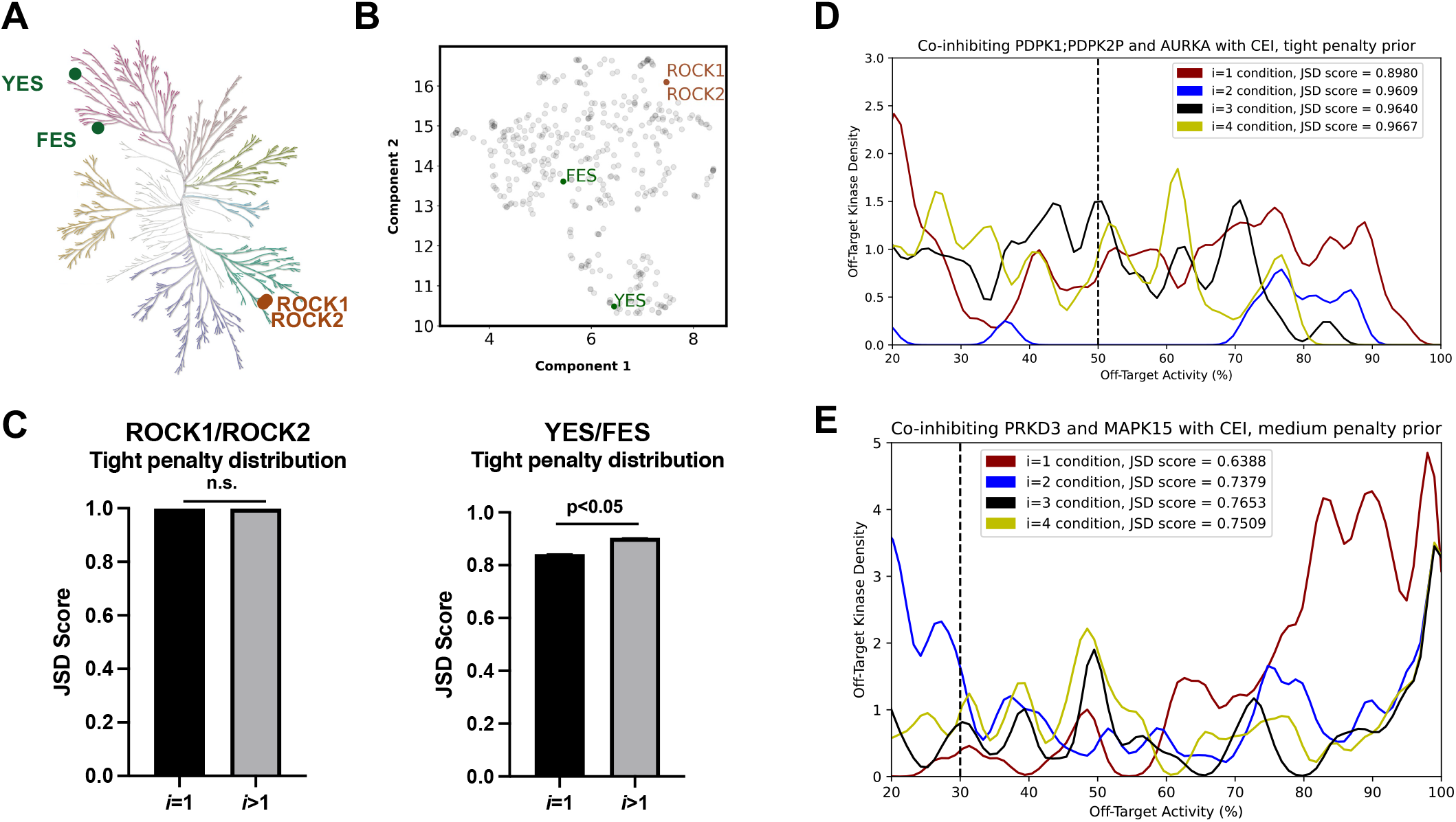
MMS identifies the most selective inhibitor set for multiple target kinases. Kinases ROCK1 and ROCK2 are closely related, while kinases YES and FES, while somewhat related, are not nearest neighbors (A). UMAP analysis of PKIS2 inhibitor phenotypes by kinase illustrates that ROCK1 and ROCK2 are inhibited by very similar inhibitors, while YES and FES are divergent in the two-component space (B). Optimization of off-target effects using PKIS2 combinations of up to three inhibitors, while maintaining activity against both members of the YES/FES or ROCK1/ROCK2 pairs at ≥ 90%, reveals that a single inhibitor performs as well as a combination for ROCK1/ROCK2, while a combination of inhibitors reduces off-target effects for YES/FES relative to the single most selective inhibitor against both kinases; data plotted as mean ± std (C). Combinations of three or four clinically evaluated inhibitors (CEI) reduce off-target effects while maintaining potent (>90%) inhibition for target kinase pairs selected from the Klaeger *et al* dataset of CEI (D, E). Significantly improved off-target profiles for target kinase pairs were assessed using the tight penalty prior (D) or medium penalty prior (E), are indicated. Dotted lines represent approximate cutoff of the penalty prior used to score off-target profiles. PRKD3/MAPK15 combinations were also significant when scored with the tight penalty prior (Table 1).

ROCK1 and ROCK2 are closely related serine/threonine kinases and exhibit virtually identical inhibitor phenotypes. Unsurprisingly, the most selective single inhibitor for both YES and FES is outperformed by a combination of inhibitors, while the most selective single inhibitor for ROCK1 and ROCK2 cannot be outperformed given the inhibitor combinations available in the dataset.

Unlike in our single-target analysis of the Klaeger *et al* dataset, in numerous cases the MMS approach outperforms the single most selective inhibitor or pair of inhibitors for dual kinase targets. We identify kinase pairs using the Klaeger *et al* dataset that are more selectively inhibited using three or four clinically evaluated inhibitors (CEI) than the single most selective inhibitor or pair of most selective CEI, or whether: *max*(*i*=3, *i*=4) > *max*(*i*=1, *i*=2). For example, PDPK1;PDPK2P, a dual-assignment made in the Klaeger *et al* dataset, and AURKA are a candidate pair identified using the tight penalty prior (Fig 6d). Phosphoinositide-dependent protein kinase-1 (PDPK1) and Aurora kinase A (AURKA) are co-targets for dual-specificity inhibitor development in the treatment of Ewing sarcoma and glioblastoma multiforme [38, 39]. The top-performing combinations of two inhibitors, SAR-407899 with either Alisertib or MLN-8054, outperforms the most selective single inhibitor, Lestaurtinib, while maintaining 90% activity against both kinases. However, the top-scoring combinations of four inhibitors: Alisertib, Lestaurtinib, SAR-407899, and either PF-03814735, XL-228, or AT-9283, outperform all combinations of two inhibitors with the tight penalty prior. Variance introduced in our calculations to compensate for measurement and calculation error indicates that there are three different inhibitors that can fill the fourth position in the *i=4* combination; calculated off-target activities for these compounds are close enough that all are identified as possibilities. These combinations of four inhibitors reduce the five highest off-target effects, calculated to be between 80% and 90% activity, to under 80% activity. Scoring was not significant when queried using the medium penalty prior: higher-order combinations did not reduce lower magnitude off-target effects.

However, 56 other kinase pairs, identified in a small non-exhaustive screen of kinase targets from the Klaeger *et al* dataset, were found to be significantly more selectively inhibited for both the tight and medium penalty priors at (*i*=3, *i*=4) than (*i*=1, *i*=2) (Table S1, Fig 6e).

Kinases were included in the analysis if their best single-inhibitor JSD score was equal to or less than 0.95. The magnitude of the improvement in JSD scores were also greater than the improvements observed for single targets from the same dataset, with many targets approaching or exceeding JSD > 0.01, a full order of magnitude greater than the 0.001 cutoff we had imposed.

These data indicate that, for target sets where highly selective inhibitors with action against all targets are not available, combinations of inhibitors that are greater than the number of targets can be calculated to minimize off-target effects while retaining high (>90%) activity against all targets. It may also be desirable to consider how to most selectively inhibit *N* clinically relevant co-target kinases, where *N* is equivalent to the number of kinases. In these cases with *N* co-targets, even if *N* inhibitors (an equal number of inhibitors to kinases) yield the highest JSD score, the most selective combination of *N* inhibitors is not necessarily the set of the most selective inhibitors for each target. The MMS approach identifies the most selective combination in these cases. Lastly, these data raise the intriguing possibility that there may be combinations of clinically evaluated inhibitors within pharmacologically reasonable combination ranges (3 or 4 inhibitors) that could more selectively inhibit kinase co-targets than the most selective compounds (1 or 2 inhibitors) identified for single kinase targets in the set. The MMS approach is extendible to >2 target kinases or higher order inhibitor combinations. Such combinations would extend the utility of current inhibitor toolsets for studying the effects of kinase inhibition in biological systems.

## Conclusion

Here, we demonstrate that we can increase the selective inhibition of a target kinase, or set of target kinases, over that of the single most selective inhibitor, even for cases where the single most selective inhibitor has very few off-target effects. JSD scoring of cumulative off-target profiles presents a new twist on an old theme: an entropy-based scoring approach to flexibly score ranges of off-target inhibitor effects, and consider ensembles of off-target profiles, that can be applied to single or multiple targets. The JSD score can also be used as a selectivity metric. Sensitivity to prior shape allows the JSD scoring approach to score off-target activity in the particular activity range that a user is most interested in. Use of complementary priors allows a user to study global effects on selectivity. Simulated data demonstrates that larger inhibitor sets, with greater combinatorial options, increases the utility of this method even as the most selective single inhibitor case also improves (Fig 3). Predictions of cumulative compound activity and inhibitor combination selectivity are well-validated using the NanoBRET target engagement assay (Fig 5). Analysis with the Klaeger *et al* dataset suggests that, while selective inhibitors may already be available for many targets, off-target effects for co-target sets can be reduced by a large magnitude and may be of greater interest (Fig 6). This framework may be of particular translational interest for polypharmacological co-inhibition of therapeutic targets, such as kinases with compensatory signaling mechanisms or those at different stages of the same signaling pathway. The MMS approach may also be extended to pharmacologic scenarios beyond the inhibition of kinases, when minimizing off-target effects are desirable and multiple interventions against the same target are available.

## Statement on code and data availability

Code, datasets, and results are available at: https://github.com/iouthwaite/inhibitor_combinations

## Author Contributions

M.A.S. conceptualized project. I.R.O., S.S., and M.A.S. designed the MMS method; I.R.O. and S.S. wrote software. I.R.O. performed MMS experiments. B.-T.B. performed NanoBRET assays. All of the authors provided feedback and discussed experiments. I.R.O. wrote the initial draft of the manuscript and all authors edited and approved the manuscript. M.A.S., J.D.C., and S.K. supervised research and acquired funding.

## Acknowledgements

M.A.S. acknowledges funding by NIH R35GM119437. I.R.O. is supported by NIH T32GM136572 and NIH T32GM008444. J.D.C. and S.S. acknowledge funding from MSKCC and NIH grant R01GM121505. S.S. is a Damon Runyon Quantitative Biology Fellow supported by the Damon Runyon Cancer Research foundation (DRQ-14-22). S.K. and B.-T.B. are grateful for support from the SGC, a registered charity (no. 1097737) that receives funds from AbbVie, Bayer, Boehringer Ingelheim, the Canada Foundation for Innovation, Eshelman Institute for Innovation, Genentech, Genome Canada through Ontario Genomics Institute (OGI-196), EU/EFPIA/OICR/McGill/KTH/Diamond, Innovative Medicines Initiative 2 Joint Undertaking (EUbOPEN grant 875510), Janssen, Merck, Merck & Co, Pfizer, Takeda and Wellcome; S.K. from the German translational cancer network (DKTK) and the Frankfurt Cancer Institute (FCI); and S.K. and B.-T.B. from the collaborative research center 1399 “Mechanisms of drug sensitivity and resistance in small cell lung cancer”. We would like to thank Dr. David Drewry and Dr. Mohammad Anwar Hossain at the Structural Genomics Consortium and Division of Chemical Biology and Medicinal Chemistry, UNC Eshelman School of Pharmacy, University of North Carolina at Chapel Hill, Chapel Hill, for providing the compounds used in this work.

