## supplement for "Death by a Thousand Cuts – Combining Kinase Inhibitors for Selective Target Inhibition and Rational Polypharmacology"

† To whom correspondence should be sent:

Markus Seeliger

### Supplementary Information

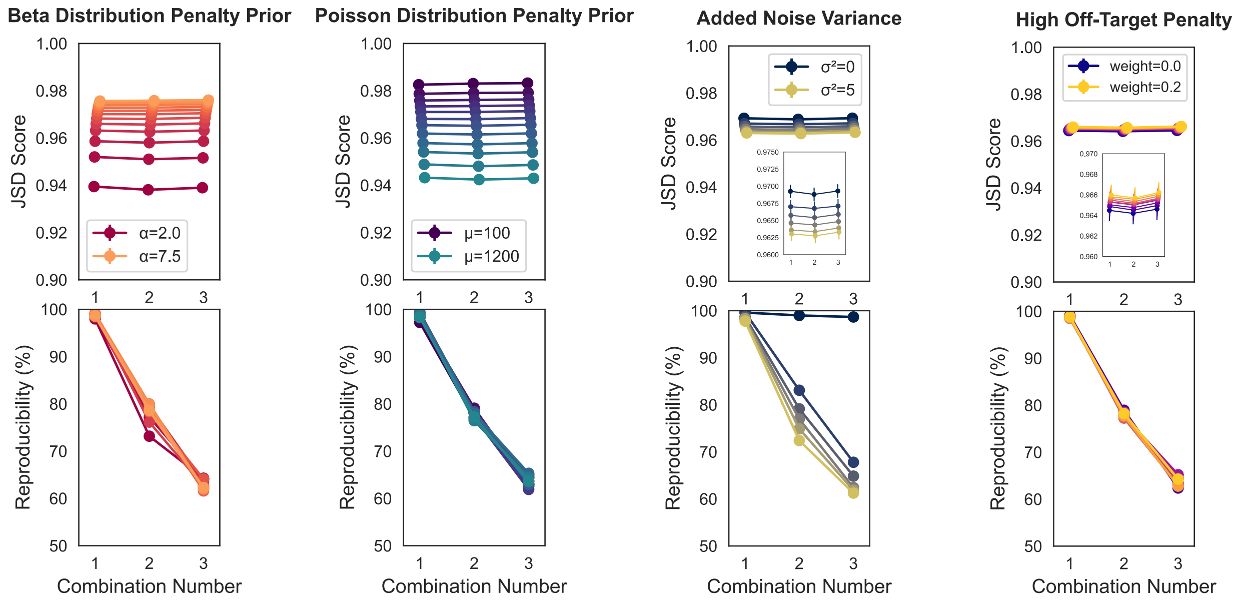


### S1. Variation of primary method parameters

Increasing the broadness of the penalty distribution, with either decreasing values of α for a beta distribution prior shape, or increasing values of μ for a Poisson distribution prior shape, increases the range of off-target effects that are penalized and correspondingly decreases the average JSD score assigned across all values of *i*. Increasing the variance of the normal distribution that noise is sampled from to build off-target distributions decreases the reproducibility of the method, with minimal impact on scoring trend. Increasing the penalty assigned to the highest off-target effects has minimal impact on scoring trends and reproducibility. Five technical replicates were performed for all analyses, using the PKIS2 dataset without promiscuous kinases, in order to increase the speed of the analysis. Data plotted as mean ± SEM. Default parameters were a Poisson medium (μ=700) penalty prior, a noise distribution variance of 2.5, and a high off-target penalty of 0.1. Reproducibility was harshly assessed as the percentage of results across the five technical replicates with identical inhibitor sets for the highest-scoring combination. For example, if across five replicates at the *i*=3 condition two of the five sets of results differed at only a single inhibitor out of three, then the reproducibility of those five replicates would be scored at 60%. Slight variation is expected with no noise due to alternate sampling of the penalty prior between replicates, as observed.

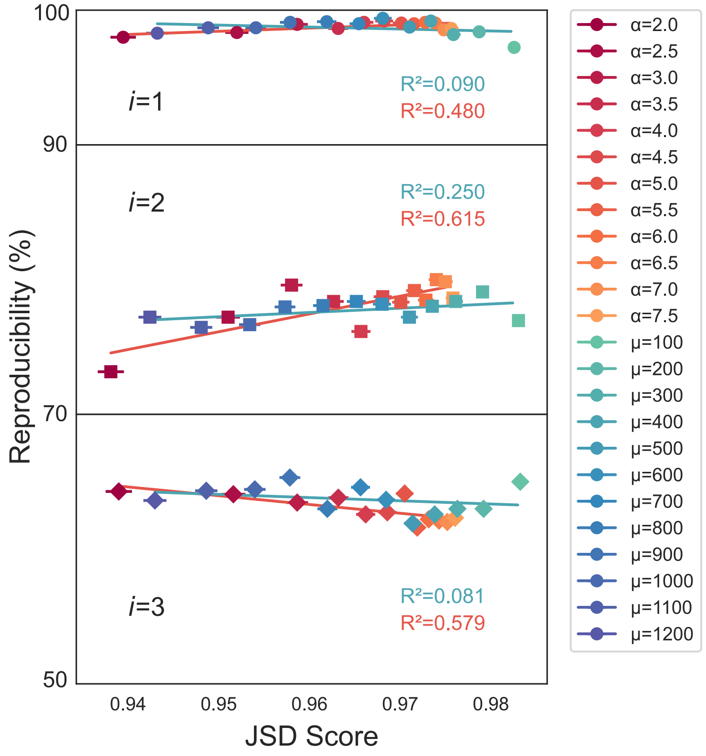


**S2. Effect of penalty prior shape on JSD score and reproducibility**

Changing the underlying shape of the penalty prior does not have an outsized impact on the reproducibility of the method, as assessed by reproducibility per JSD score. The reproducibility of scores generated using the Poisson distribution penalty prior is not associated with the assigned score. For this reason, the Poisson distribution penalty prior was selected for primary method analyses.

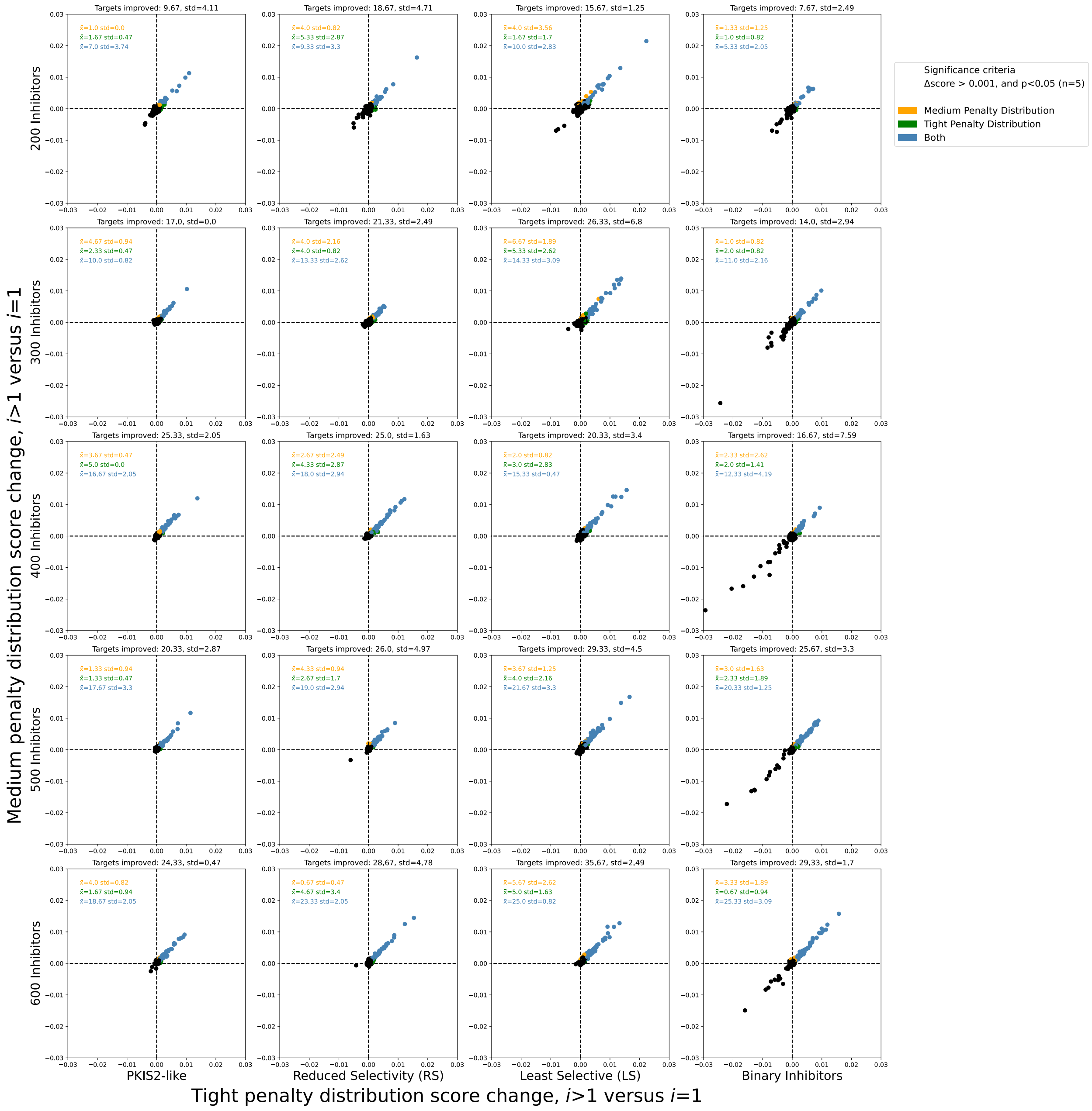


**S3. Simulated data suggests improvement in off-target effects using inhibitor combinations, with variation in inhibitor profiles and inhibitor set size: figure 3 extended data**

Simulated data generated using inhibitor sets (Fig 3) was analyzed using both a tight penalty distribution and a medium penalty distribution. All targets across three experimental replicates of each condition (300) are plotted; the mean ± std of significantly improved targets across these three replicates, for each penalty prior and both, are indicated. Significance was assessed by statistical significance across five technical replicates for each experimental replicate, and an absolute average score increase of at least 0.001.

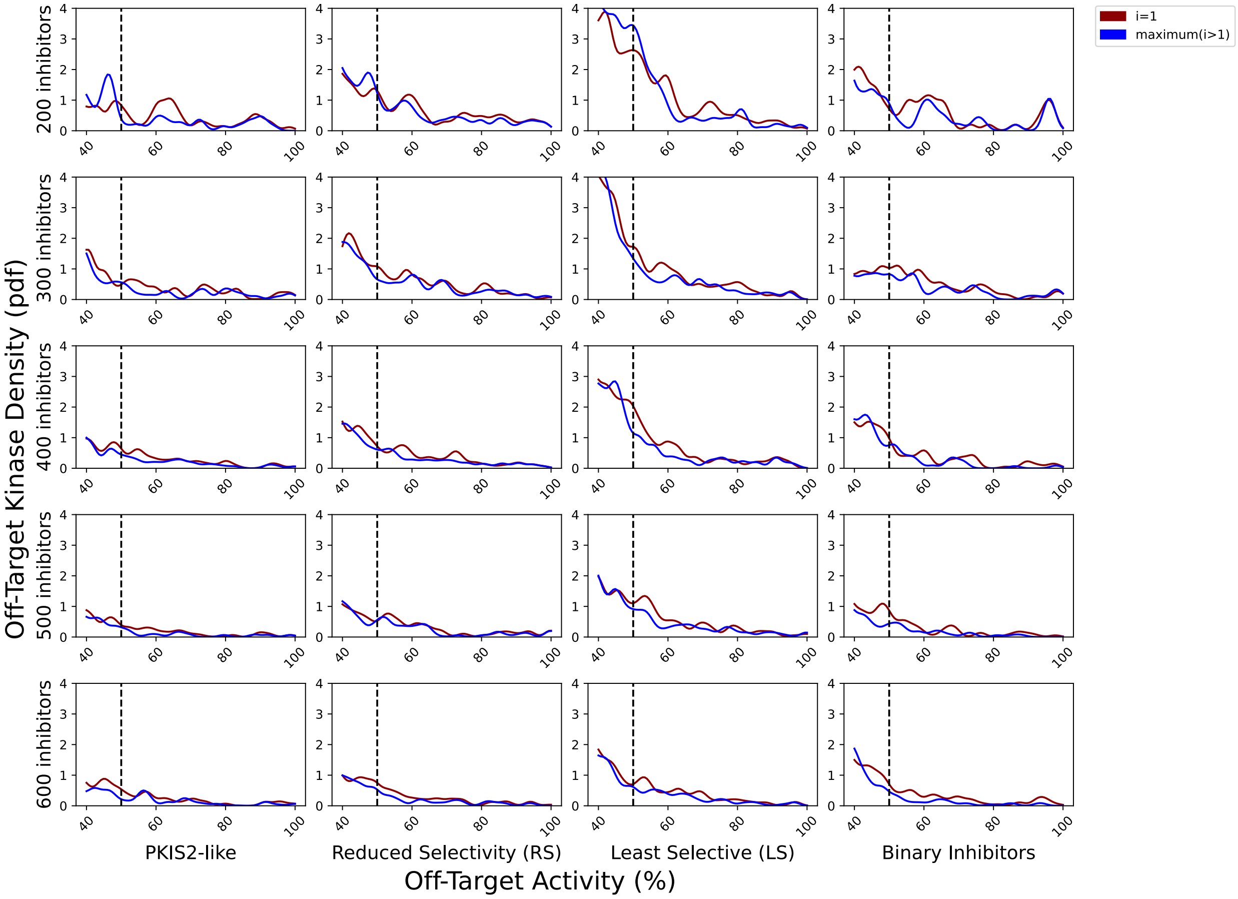


**S4. Simulated data off-target activity profiles using a tight penalty distribution and a threshold of 0.001 suggest improvements in off-target effects using inhibitor combinations: figure 3 extended data**

Probability density functions illustrate the average off-target effects for the highest scoring single inhibitor versus the highest scoring combination of inhibitors for simulated datasets, scored using a tight penalty distribution with an absolute JSD score increase of ≥0.001. Vertical dotted lines represent the approximate activity value where the tight probability distribution becomes negligible. The variance of the normal distributions used to generate the pdfs is the same as was used in the scoring method: 2.5, such that the overall density is an accurate reflection of the average off-target distributions.

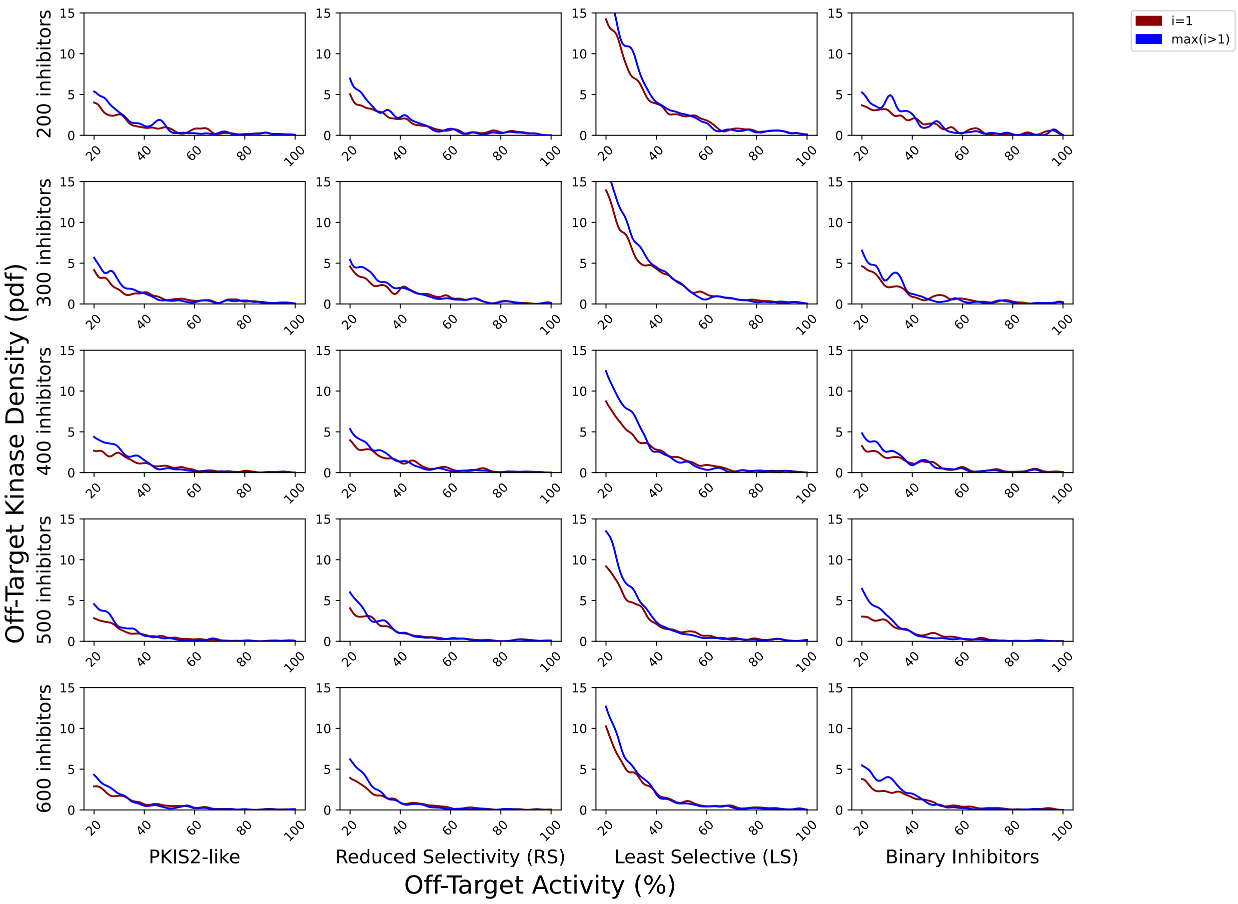


**S5. Simulated data off-target activity profiles using a medium penalty distribution and a threshold of 0.001 suggest improvements in off-target effects using inhibitor combinations: figure 3 extended data**

Probability density functions illustrate the average off-target effects for the highest scoring single inhibitor versus the highest scoring combination of inhibitors for simulated datasets, scored using a medium penalty distribution with an absolute JSD score increase of ≥0.001. The variance of the normal distributions used to generate the pdfs is the same as was used in the scoring method: 2.5, such that the overall density is an accurate reflection of the average off-target distributions. Like the other penalty distributions, the medium off-target penalty distribution penalizes higher off-target effects much more than lower off-target effects, and becomes negligible at approximately 30% activity.

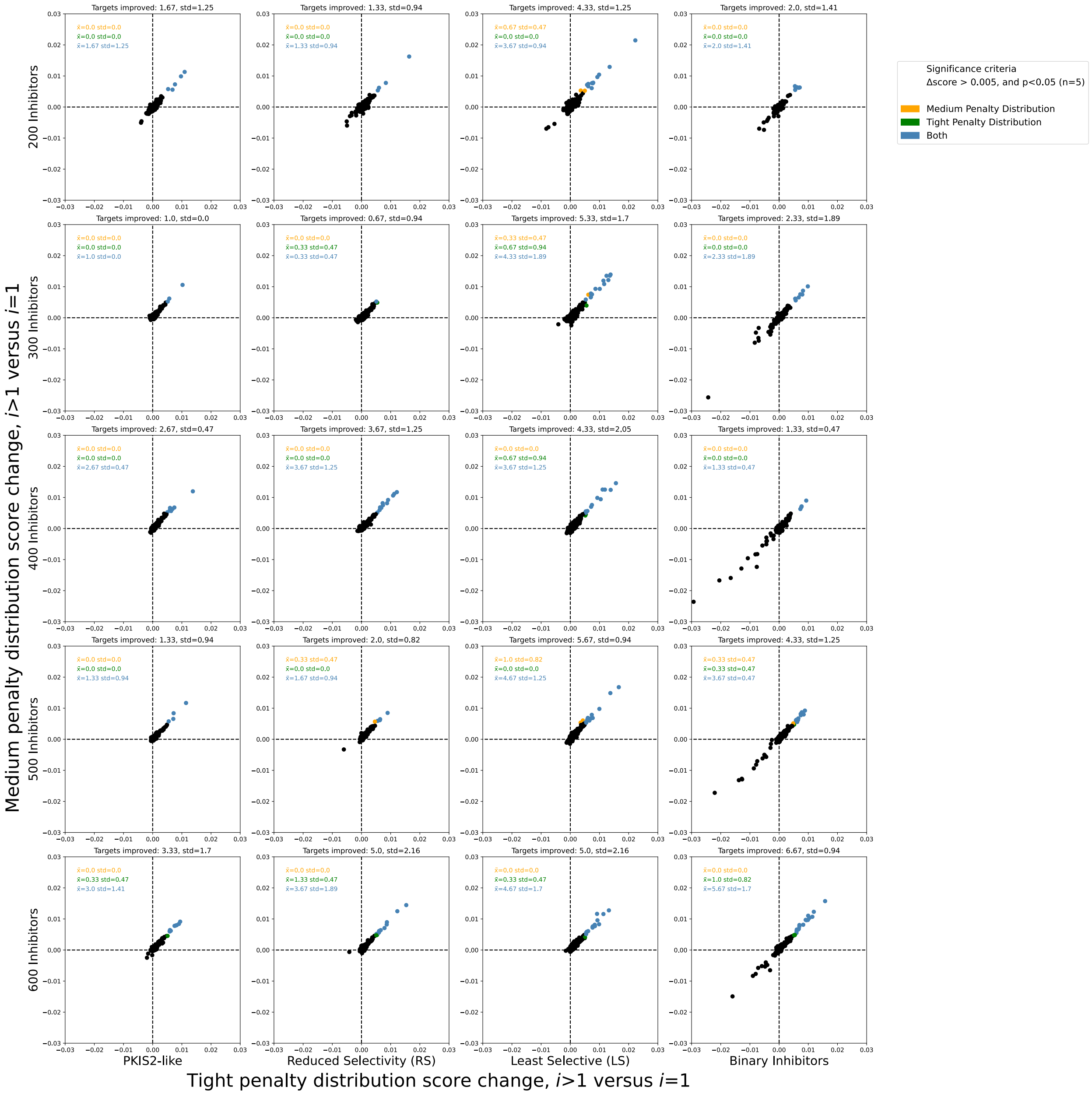

**S6. Increasing the absolute JSD score cutoff to 0.005 decreases observed off-target effects: figure 3 extended data**

Simulated data generated using inhibitor sets (Fig 3) was analyzed using both a tight penalty distribution and a medium penalty distribution. All targets across three experimental replicates of each condition (300) are plotted; the mean ± std of significantly improved targets across these three replicates, for each penalty prior and both, are indicated. Significance was assessed by statistical significance across five technical replicates for each experimental replicate, and an absolute average score increase of at least 0.005.

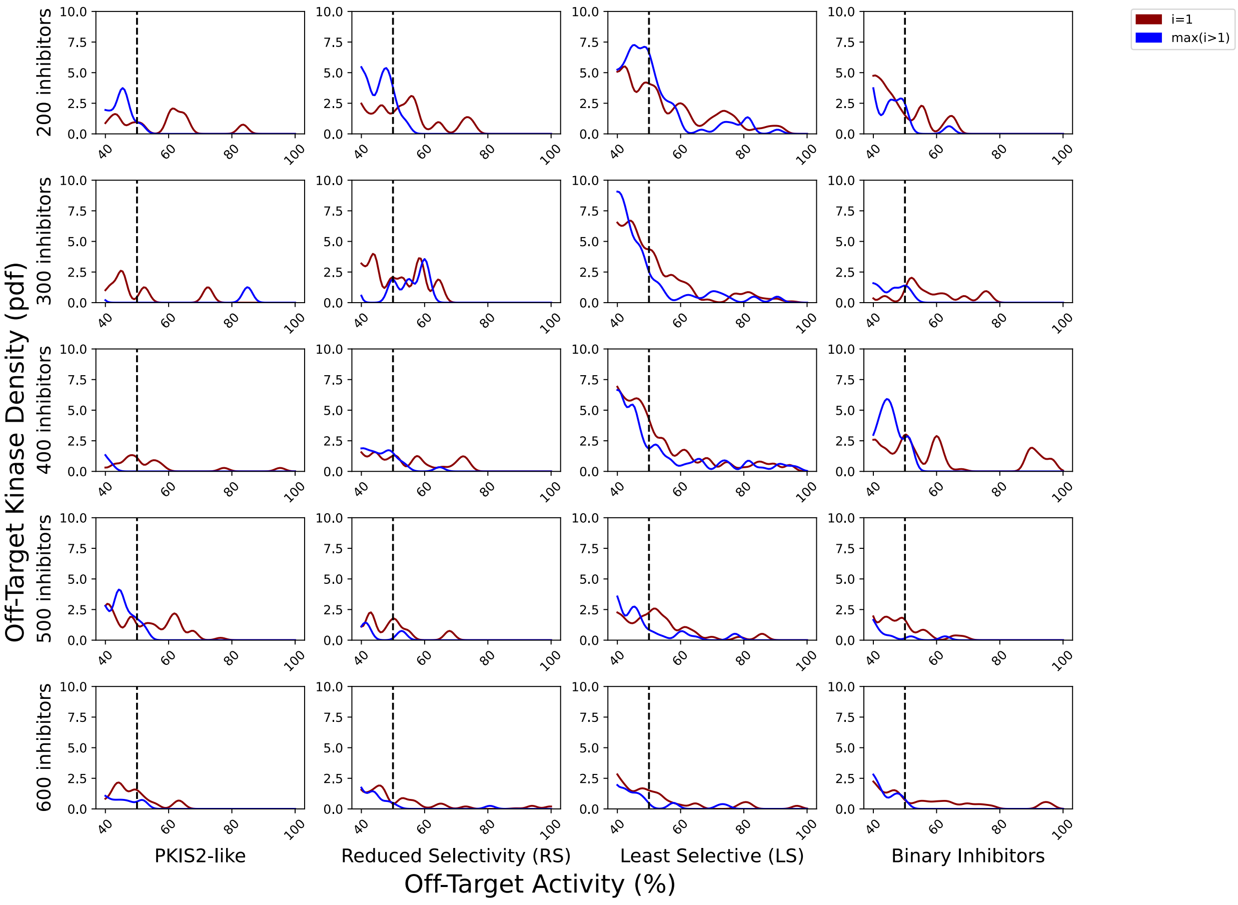

**S7. Simulated data off-target activity profiles using a tight penalty distribution and a threshold of 0.005 suggest greater improvements in off-target effects using inhibitor combinations: figure 3 extended data**

Probability density functions illustrate the average off-target effects for the highest scoring single inhibitor versus the highest scoring combination of inhibitors for simulated datasets, scored using a tight penalty distribution with an absolute JSD score increase of ≥0.005. Vertical dotted lines represent the approximate activity value where the tight probability distribution becomes negligible. The variance of the normal distributions used to generate the pdfs is the same as was used in the scoring method: 2.5, such that the overall density is an accurate reflection of the average off-target distributions.

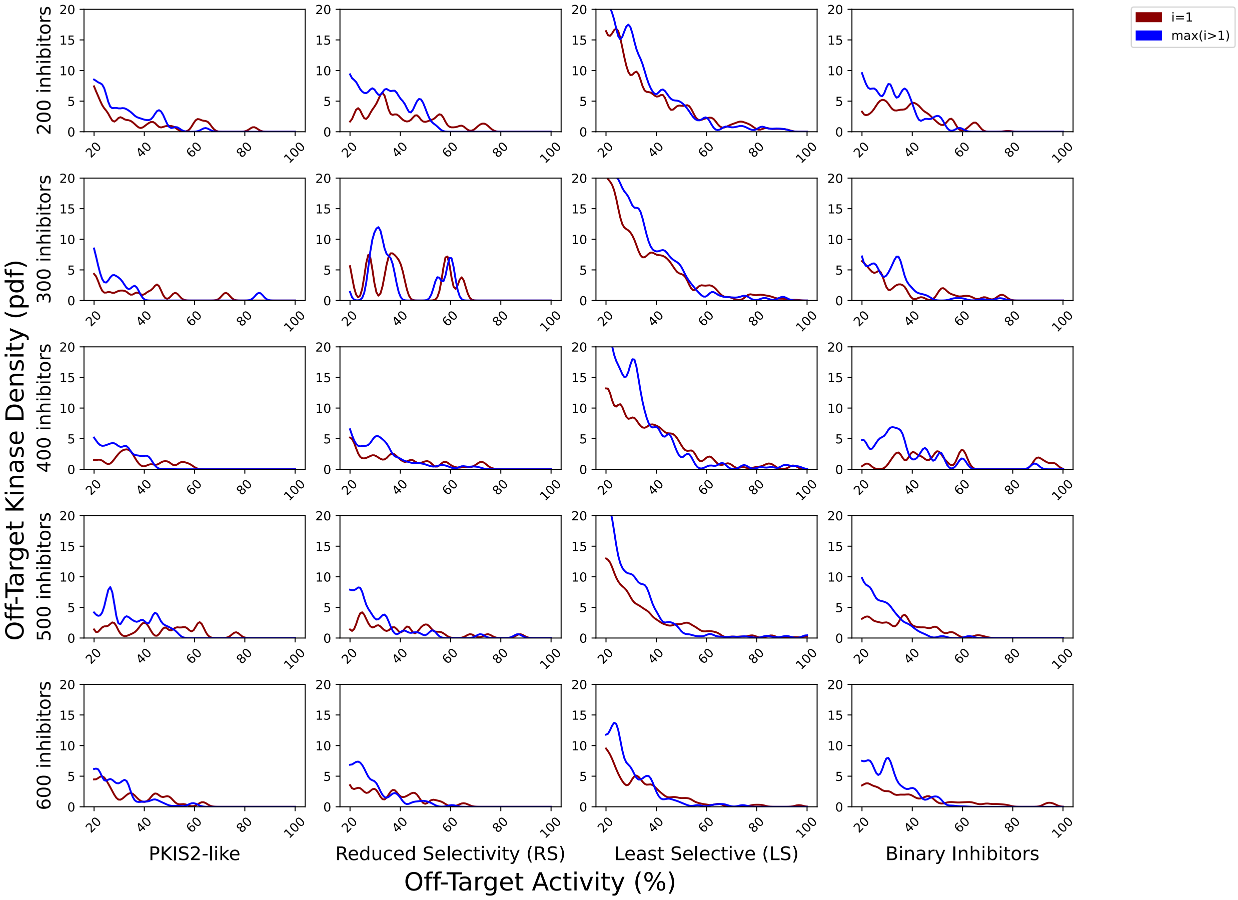


**S8. Simulated data off-target activity profiles using a medium penalty distribution and a threshold of 0.005 suggest greater improvements in off-target effects using inhibitor combinations: figure 3 extended data**

Probability density functions illustrate the average off-target effects for the highest scoring single inhibitor versus the highest scoring combination of inhibitors for simulated datasets, scored using a medium penalty distribution with an absolute JSD score increase of ≥0.005. Vertical dotted lines represent the approximate activity value where the tight probability distribution becomes negligible. The variance of the normal distributions used to generate the pdfs is the same as was used in the scoring method: 2.5, such that the overall density is an accurate reflection of the average off-target distributions.

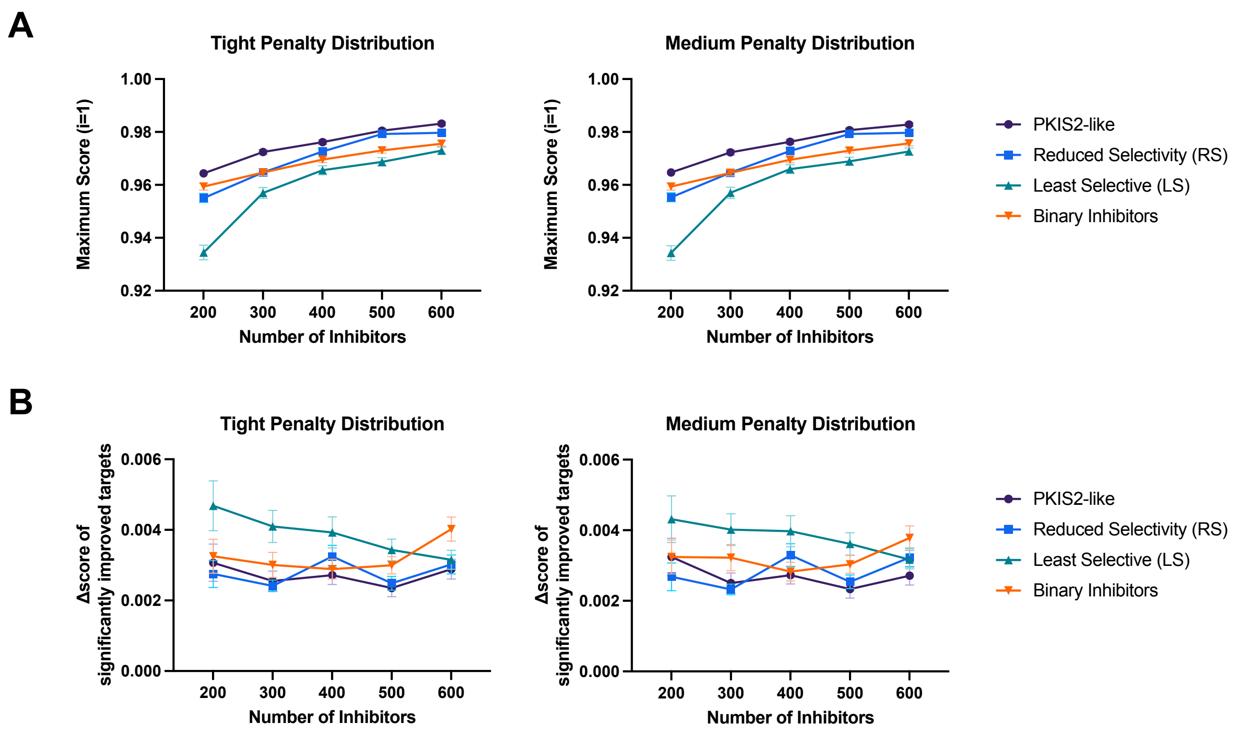


**S9. Increasing inhibitor dataset coverage improves inhibitor selectivity for all values of *i:* figure 3 extended data**

Increasing the number of simulated inhibitors, or the fold-coverage of 100 simulated targets, improved the average selectivity of the single most selective inhibitor in all cases (A). The top-scoring single inhibitors in all simulated datasets across all targets has, on average, a very similar JSD score when scored with the tight and medium penalty distributions (A). The average increase in score for significantly improved targets does not appear to change for PKIS2-like or slightly Less Selective B inhibitor combinations (B). The average increase in score tends to decrease for Least Selective (LS) inhibitor combination types, as the average score of the single most selective inhibitor increases and approaches selectivity closer to that of PKIS2-like or Reduced Selectivity (RS) type inhibitors (B). Combinations of simulated binary inhibitors may trend towards higher magnitude of performance with increasing fold-coverage of targets, but this is limited due to the maximum screened set size and may be influenced by an outlier. Data plotted as mean ± SEM for three experimental condition replicates, with significance assessed by five technical replicates ($p < 0.05$, absolute score change > 0.001) for each of the three experimental condition replicates.

| Rank | Kinase | ΔJSD μ=200 | ΔJSD μ=700 | ΔJSD μ=200 stdev | ΔJSD μ=700 stdev | i=1 score μ=200 | i=1 score μ=200 stdev | i>1 max score μ=200 | i>1 max score μ=200 stdev | i=1 score μ=700 | i=1 score μ=700 stdev | i>1 max score μ=700 | i>1 max score μ=700 stdev |
|---|---|---|---|---|---|---|---|---|---|---|---|---|---|
| 1 | PLK3 | 0.00524 | 0.011419 | 0.000108 | 0.000613 | 0.981691 | 0.000129 | 0.986932 | 6.3e-05 | 0.935531 | 0.000745 | 0.94695 | 0.000871 |
| 2 | p38-alpha | 0.004321 | 0.009692 | 7.7e-05 | 0.000194 | 0.99421 | 8.9e-05 | 0.998531 | 6.3e-05 | 0.966733 | 0.000636 | 0.976424 | 0.000561 |
| 3 | RIPK2 | 0.004702 | 0.005581 | 0.000115 | 0.001279 | 0.976469 | 9.2e-05 | 0.981171 | 7.7e-05 | 0.92563 | 0.000329 | 0.931211 | 0.001494 |
| 4 | PLK2 | 0.002284 | 0.006993 | 6.9e-05 | 0.000682 | 0.986013 | 0.000132 | 0.988296 | 8.6e-05 | 0.94743 | 0.000931 | 0.954423 | 0.000658 |
| 5 | p38-beta | 0.001958 | 0.005125 | 0.000121 | 0.00058 | 0.991274 | 0.000183 | 0.993232 | 0.000121 | 0.950566 | 0.000696 | 0.955691 | 0.001124 |
| 6 | JNK2 | 0.002646 | 0.002041 | 0.000183 | 0.000229 | 0.987218 | 0.000181 | 0.989864 | 0.000151 | 0.969261 | 0.00051 | 0.971303 | 0.000588 |
| 7 | JNK1 | 0.002795 | 0.001817 | 9.4e-05 | 0.000894 | 0.987161 | 0.000129 | 0.989957 | 0.000116 | 0.969403 | 0.000338 | 0.97122 | 0.000749 |
| 8 | ACVR1 | 0.001165 | 0.003369 | 9.7e-05 | 5.2e-05 | 0.993047 | 0.000127 | 0.994212 | 4.1e-05 | 0.967785 | 0.000642 | 0.971154 | 0.000596 |
| 9 | LIMK1 | 0.001182 | 0.003227 | 7e-05 | 9.4e-05 | 0.993013 | 0.000145 | 0.994195 | 8.4e-05 | 0.967591 | 0.000743 | 0.970818 | 0.000824 |
| 10 | JNK3 | 0.002667 | 0.001686 | 6.9e-05 | 0.000333 | 0.987173 | 0.000111 | 0.98984 | 7.8e-05 | 0.969527 | 0.000518 | 0.971212 | 0.000622 |
| 11 | ZAK | -0.000603 | 0.004946 | 0.000129 | 0.000554 | 0.985886 | 0.000111 | 0.985283 | 8.6e-05 | 0.952018 | 0.000611 | 0.956964 | 0.000704 |
| 12 | TXK | -0.000686 | 0.004969 | 7.1e-05 | 0.000238 | 0.985995 | 9.7e-05 | 0.985309 | 7.4e-05 | 0.951862 | 0.000644 | 0.956831 | 0.000695 |
| 13 | TESK1 | -0.000502 | 0.004192 | 0.00011 | 0.000888 | 0.986177 | 0.000211 | 0.985675 | 0.000139 | 0.952573 | 0.000671 | 0.956765 | 0.000939 |
| 14 | KIT | 0.000549 | 0.002786 | 0.000129 | 0.000464 | 0.977986 | 0.000131 | 0.978535 | 7.5e-05 | 0.943257 | 0.000389 | 0.946044 | 0.000654 |
| 15 | LKB1 | -0.001214 | 0.004488 | 0.000116 | 0.000806 | 0.979765 | 0.000313 | 0.978551 | 0.000232 | 0.929783 | 0.001614 | 0.934271 | 0.000877 |
| 16 | VRK2 | 0.003052 | -2.7e-05 | 0.000365 | 0.000761 | 0.988382 | 8.6e-05 | 0.991434 | 0.000307 | 0.95041 | 0.000823 | 0.950383 | 0.000632 |
| 17 | PIK3CB | 0.002273 | 0.000584 | 7e-05 | 0.000747 | 0.996156 | 0.000146 | 0.998429 | 9.8e-05 | 0.973236 | 0.000511 | 0.973819 | 0.000777 |
| 18 | FGFR2 | -0.002674 | 0.003735 | 0.00011 | 0.000157 | 0.981404 | 0.000165 | 0.97873 | 0.000155 | 0.934192 | 0.001176 | 0.937927 | 0.001254 |
| 19 | ROCK1 | 0.001832 | -0.001848 | 0.000172 | 0.000903 | 0.989719 | 0.000117 | 0.991551 | 9.9e-05 | 0.954122 | 0.000595 | 0.952274 | 0.000821 |
| 20 | PLK1 | -0.003588 | 0.003105 | 0.000155 | 0.001032 | 0.992589 | 0.000152 | 0.989001 | 0.0001 | 0.955083 | 0.001331 | 0.958188 | 0.000639 |
| 21 | MARK4 | -0.006496 | 0.005235 | 9.4e-05 | 0.000511 | 0.988104 | 0.000178 | 0.981608 | 0.000221 | 0.957957 | 0.000478 | 0.963192 | 0.00064 |
| 22 | MARK1 | -0.006514 | 0.004939 | 0.000109 | 0.000206 | 0.988097 | 0.000182 | 0.981583 | 0.000226 | 0.958234 | 0.000694 | 0.963174 | 0.000644 |
| 23 | BRSK2 | -0.006535 | 0.004738 | 0.000131 | 0.000117 | 0.988097 | 0.000101 | 0.981562 | 8.8e-05 | 0.958808 | 0.000532 | 0.963546 | 0.000581 |
| 24 | TNNI3K | -0.009223 | 0.002247 | 0.000239 | 0.000702 | 0.976479 | 0.000123 | 0.967255 | 0.000134 | 0.918402 | 0.000702 | 0.920649 | 0.00099 |

### S10. Kinases with significant improvements in off-target effects using combinations of PKIS2 inhibitors: figure 4 extended data

Kinase targets were assessed using combinations of inhibitors from PKIS2. Tight penalty distribution and medium penalty distribution priors were used (see main text Fig 4). Five technical replicates were performed for all analyses with significance for each penalty distribution prior assessed by $p < 0.05$, and an absolute score change > 0.001. Noise distribution variance was set at 2.5, and high on-target penalty set at 0.1.

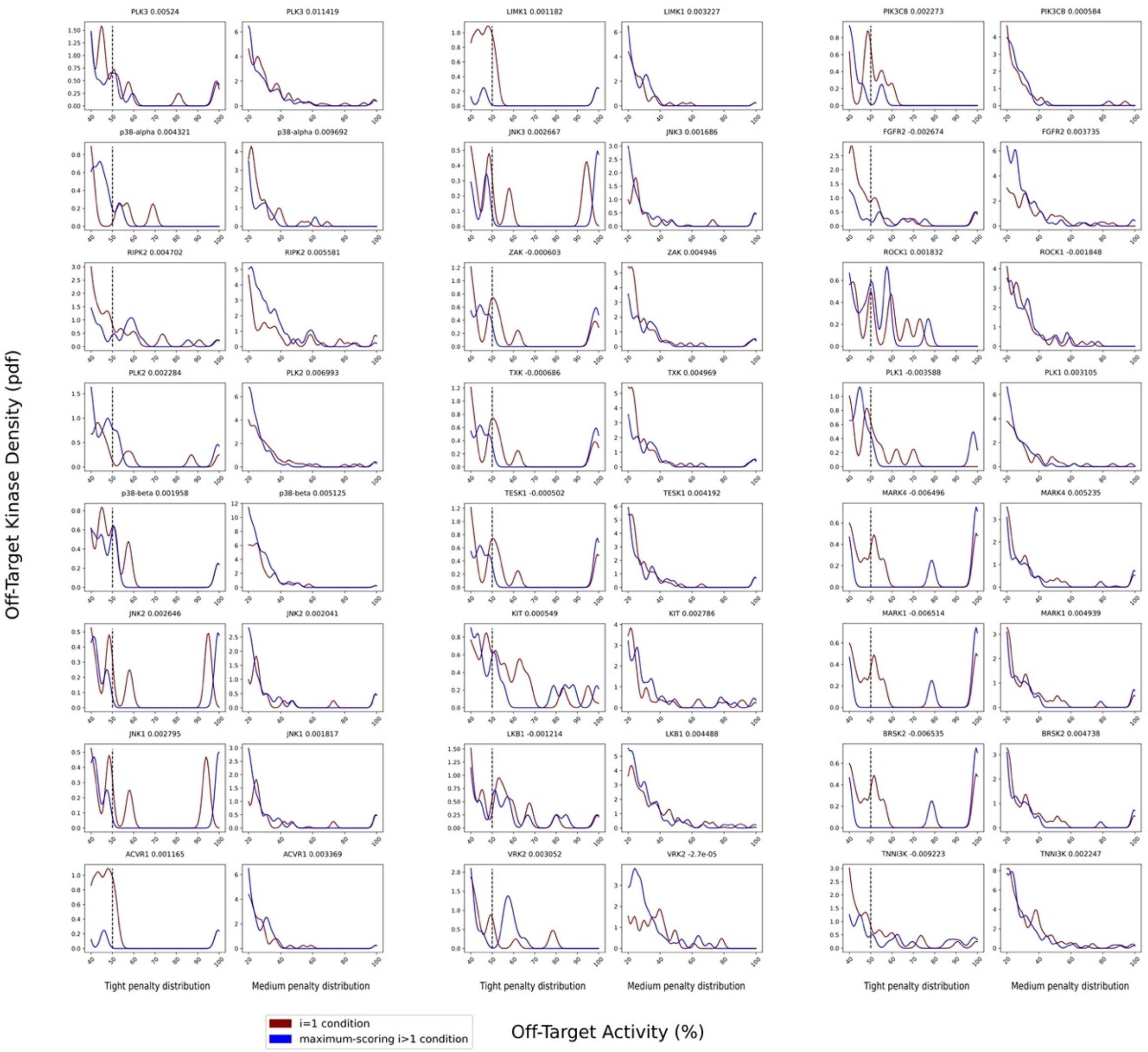


**S11. Off-target activity profiles suggest improvements in off-target effects using inhibitor combinations with inhibitors from PKIS2: figure 4 extended data**

Probability density functions illustrate the average off-target effects for the highest scoring single inhibitor versus the highest scoring combination of inhibitors for inhibitor combinations from PKIS2, scored using either a tight or medium penalty distribution with an absolute JSD score increase of >=0.001. The variance of the normal distributions used to generate the pdfs is the same as was used in the scoring method: 2.5, such that the overall density is an accurate reflection of the off-target penalty distributions. Vertical dotted lines represent the approximate activity value where the tight probability distribution becomes negligible. Like the tight penalty distribution, the medium penalty distribution penalizes higher off-target effects much more than lower off-target effects, and becomes negligible at approximately 30% activity. Although off-target effects for a kinase may have only been significantly improved using one of the two penalty distributions (tight or medium), data from both is presented to illustrate corresponding non-significant results.

| Rank | Kinase | ΔJSD μ=200 | ΔJSD μ=700 | ΔJSD μ=200 stdev | ΔJSD μ=700 stdev | i=1 score μ=200 | i=1 score μ=200 stdev | i>1 max score μ=200 | i>1 max score μ=200 stdev | i=1 score μ=700 | i=1 score μ=700 stdev | i>1 max score μ=700 | i>1 max score μ=700 stdev |
|---|---|---|---|---|---|---|---|---|---|---|---|---|---|
| 1 | TLK1 | 0.006188 | 0.00645 | 0.005262 | 0.00509 | 0.482844 | 0.002353 | 0.489032 | 0.003074 | 0.404253 | 0.001954 | 0.410703 | 0.004115 |
| 2 | FLT3(D835Y) | 0.002727 | 0.005686 | 0.000285 | 0.000461 | 0.992347 | 0.000302 | 0.995073 | 9.5e-05 | 0.960801 | 0.00049 | 0.966488 | 0.000817 |
| 3 | FLT3(D835H) | 0.00583 | 0.000296 | 0.000311 | 0.000164 | 0.982036 | 0.00041 | 0.987866 | 0.000327 | 0.962083 | 0.000199 | 0.962379 | 0.000321 |
| 4 | ABL1-nonphosphorylated | 0.004162 | 0.000321 | 0.000107 | 0.000242 | 0.992997 | 0.000121 | 0.997159 | 0.000188 | 0.97861 | 0.00042 | 0.978931 | 0.000505 |
| 5 | STK16 | 0.003714 | 0.000257 | 0.00078 | 0.000964 | 0.951122 | 0.000677 | 0.954836 | 0.000697 | 0.897854 | 0.000902 | 0.89811 | 0.001137 |
| 6 | FLT3(ITD) | 0.003733 | 0.000134 | 0.000393 | 0.000204 | 0.984214 | 0.000383 | 0.987947 | 0.00031 | 0.972932 | 0.0003 | 0.973066 | 0.000206 |
| 7 | FLT3 | 0.002205 | 0.000294 | 2.8e-05 | 0.000269 | 0.992898 | 9.7e-05 | 0.995103 | 0.000117 | 0.979434 | 0.000308 | 0.979729 | 0.000178 |
| 8 | FLT4 | 0.001307 | 0.001148 | 0.000293 | 0.000414 | 0.962066 | 0.00023 | 0.963373 | 0.000242 | 0.949232 | 0.000454 | 0.95038 | 0.000554 |
| 9 | FLT3(N841I) | 0.001426 | 0.001015 | 0.000388 | 0.000403 | 0.974744 | 0.000492 | 0.97617 | 0.000178 | 0.963508 | 0.000474 | 0.964522 | 0.000504 |
| 10 | GAK | 0.002319 | 0.000109 | 0.000151 | 0.000315 | 0.99141 | 7.8e-05 | 0.993729 | 8.1e-05 | 0.980753 | 0.00042 | 0.980862 | 0.000624 |
| 11 | SLK | 0.000362 | 0.002036 | 0.000524 | 0.00054 | 0.980806 | 0.000581 | 0.981168 | 0.000381 | 0.950614 | 0.000911 | 0.95265 | 0.000542 |
| 12 | ROS1 | 0.001833 | 0.000474 | 0.000178 | 0.000781 | 0.964562 | 0.00038 | 0.966395 | 0.00032 | 0.932064 | 0.000409 | 0.932538 | 0.000951 |
| 13 | ERBB4 | 3e-06 | 0.00208 | 0.000109 | 0.000193 | 0.992047 | 0.000191 | 0.99205 | 8.5e-05 | 0.982442 | 0.000212 | 0.984522 | 0.000326 |
| 14 | FLT3(K663Q) | 0.001851 | 0.000192 | 0.000208 | 0.000258 | 0.989927 | 0.000276 | 0.991778 | 0.000135 | 0.977095 | 0.000367 | 0.977287 | 0.000252 |
| 15 | ABL1(F317L)-nonphosphorylated | 0.001435 | 0.000274 | 0.000176 | 0.000237 | 0.993028 | 0.000124 | 0.994463 | 0.000205 | 0.970135 | 0.000453 | 0.970409 | 0.000538 |
| 16 | DDR1 | 6.4e-05 | 0.001066 | 3.2e-05 | 0.000261 | 0.999923 | 2.4e-05 | 0.999987 | 1.2e-05 | 0.989571 | 0.000336 | 0.990637 | 0.000438 |
| 17 | TLK2 | -0.006738 | 0.006173 | 0.000491 | 0.002218 | 0.527017 | 0.000968 | 0.520279 | 0.000842 | 0.430783 | 0.001701 | 0.436956 | 0.001906 |

### S12. Kinases with significant improvements in off-target effects using combinations of inhibitors from Davis *et al* (2011): figure 4 extended data

Kinase targets were assessed using combinations of inhibitors from Davis *et al* (2011). Tight penalty distribution and medium penalty distribution priors were used (see main text Fig 4). Five technical replicates were performed for all analyses with significance for each penalty distribution prior assessed by $p < 0.05$, and an absolute score change > 0.001. Noise distribution variance was set at 2.5, and high on-target penalty set at 0.1.

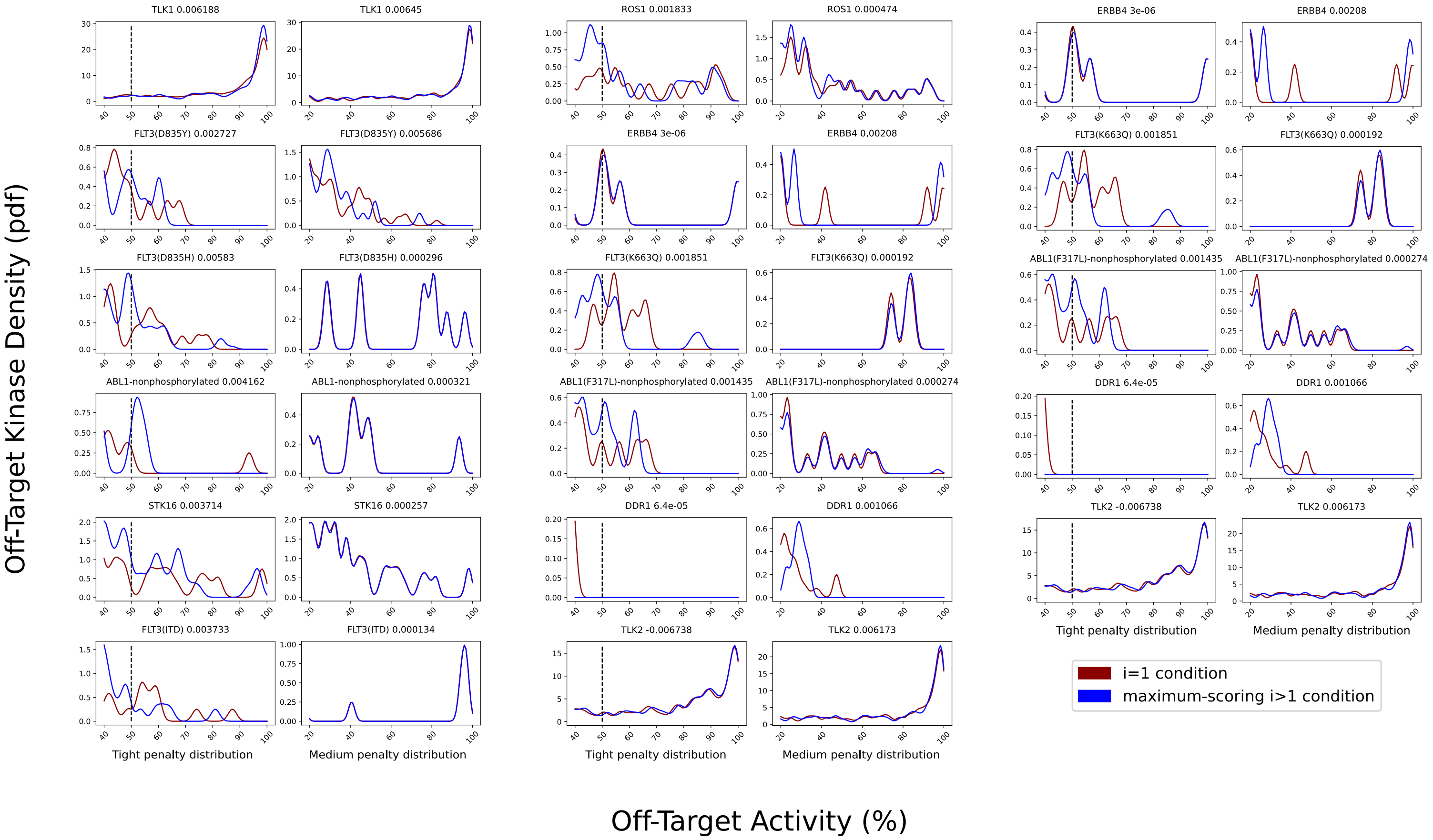


**S13. Off-target activity profiles suggest improvements in off-target effects using inhibitor combinations with inhibitors from Davis *et al* (2011): figure 4 extended data**

Probability density functions illustrate the average off-target effects for the highest scoring single inhibitor versus the highest scoring combination of inhibitors for inhibitor combinations from the Davis *et al* (2011) dataset, scored using either a tight or medium penalty distribution with an absolute JSD score increase of >=0.001. The variance of the normal distributions used to generate the pdfs is the same as was used in the scoring method: 2.5, such that the overall density is an accurate reflection of the off-target penalty distributions. Vertical dotted lines represent the approximate activity value where the tight probability distribution becomes negligible. Like the tight penalty distribution, the medium penalty distribution penalizes higher off-target effects much more than lower off-target effects, and becomes negligible at approximately 30% activity. Although off-target effects for a kinase may have only been significantly improved using one of the two penalty distributions (tight or medium), data from both is presented to illustrate corresponding non-significant results.

| Rank | Kinase | ΔJSD μ=200 | ΔJSD μ=700 | ΔJSD μ=200 stdev | ΔJSD μ=700 stdev | i=1 score μ=200 | i=1 score μ=200 stdev | i>1 max score μ=200 | i>1 max score μ=200 stdev | i=1 score μ=700 | i=1 score μ=700 stdev | i>1 max score μ=700 | i>1 max score μ=700 stdev |
|---|---|---|---|---|---|---|---|---|---|---|---|---|---|
| 1 | SLK | 0.001227 | 0.002272 | 7.6e-05 | 0.000278 | 0.978 | 6.2e-05 | 0.979227 | 5.7e-05 | 0.950441 | 0.000349 | 0.952713 | 0.000552 |
| 2 | BIKE | 0.00305 | 9.9e-05 | 0.000753 | 0.000764 | 0.88054 | 0.001205 | 0.883591 | 0.001391 | 0.807695 | 0.000846 | 0.807795 | 0.000659 |
| 3 | VEGFR2 | 0.002386 | 0.000708 | 0.000256 | 0.000295 | 0.96262 | 0.000168 | 0.965006 | 0.000134 | 0.956567 | 0.000325 | 0.957274 | 0.000158 |
| 4 | KIT(V559D,V654A) | 0.000252 | 0.002604 | 0.000237 | 0.000343 | 0.976806 | 0.000317 | 0.977058 | 0.000262 | 0.963847 | 0.000248 | 0.96645 | 0.000386 |
| 5 | JNK3 | -0.00015 | 0.001422 | 0.000469 | 0.000504 | 0.95401 | 0.000374 | 0.953861 | 0.000532 | 0.930306 | 0.000399 | 0.931728 | 0.000424 |
| 6 | FLT4 | -8.8e-05 | 0.001335 | 0.000104 | 0.000387 | 0.94859 | 0.000151 | 0.948502 | 7.8e-05 | 0.934264 | 0.000492 | 0.935598 | 0.000524 |

### S14. Kinases with significant improvements in off-target effects using combinations of inhibitors from Karaman *et al* (2008): figure 4 extended data

Kinase targets were assessed using combinations of inhibitors from Karaman *et al* (2008). Tight penalty distribution and medium penalty distribution priors were used (see main text Fig 4). Five technical replicates were performed for all analyses with significance for each penalty distribution prior assessed by $p < 0.05$, and an absolute score change > 0.001. Noise distribution variance was set at 2.5, and high on-target penalty set at 0.1.

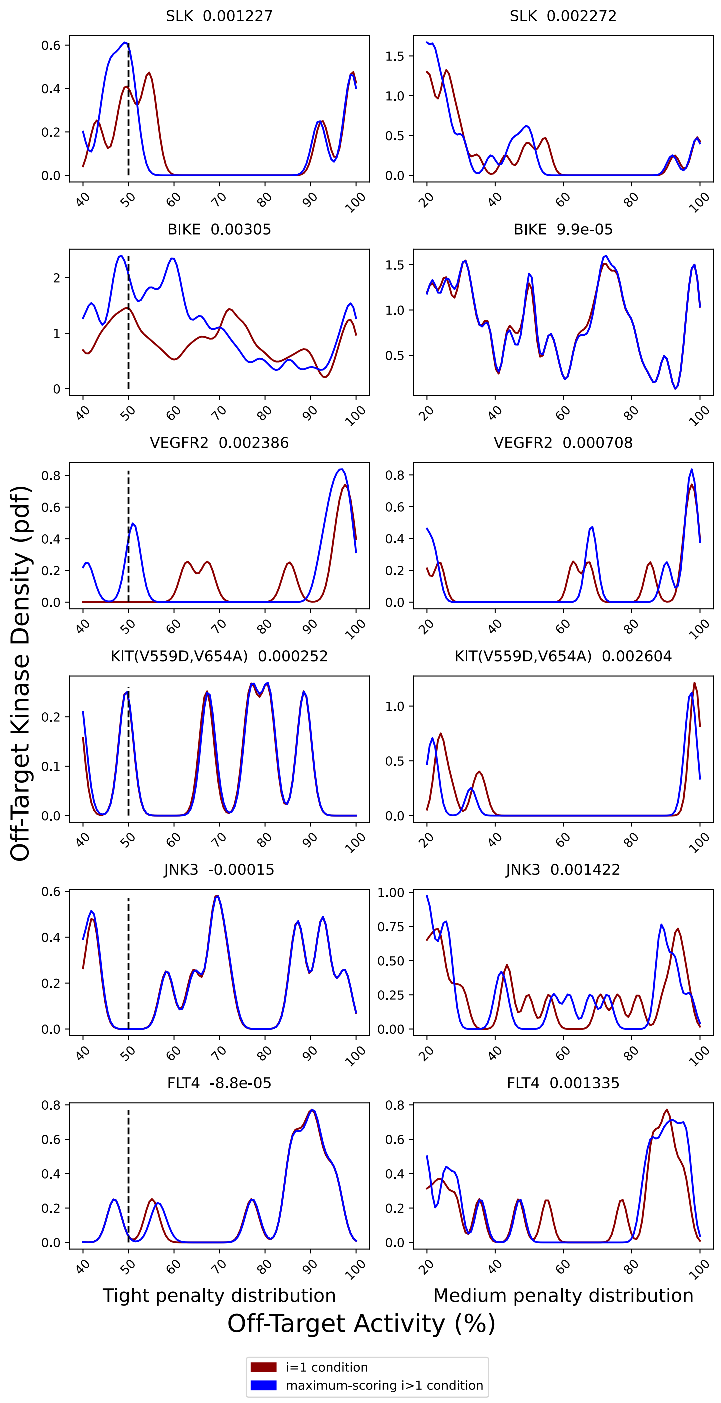


**S15. Off-target activity profiles suggest improvements in off-target effects using inhibitor combinations with inhibitors from Karaman *et al* (2008): figure 4 extended data**

Probability density functions illustrate the average off-target effects for the highest scoring single inhibitor versus the highest scoring combination of inhibitors for inhibitor combinations from the Karaman *et al* (2008) dataset, scored using either a tight or medium penalty distribution with an absolute JSD score increase of >=0.001. The variance of the normal distributions used to generate the pdfs is the same as was used in the scoring method: 2.5, such that the overall density is an accurate reflection of the off-target penalty distributions. Vertical dotted lines represent the approximate activity value where the tight probability distribution becomes negligible. Like the tight penalty distribution, the medium penalty distribution penalizes higher off-target effects much more than lower off-target effects, and becomes negligible at approximately 30% activity. Although off-target effects for a kinase may have only been significantly improved using one of the two penalty distributions (tight or medium), data from both is presented to illustrate corresponding non-significant results.

| Rank | Kinase | ΔJSD μ=200 | ΔJSD μ=700 | ΔJSD μ=200 stdev | ΔJSD μ=700 stdev | i=1 score μ=200 | i=1 score μ=200 stdev | i>1 max score μ=200 | i>1 max score μ=200 stdev | i=1 score μ=700 | i=1 score μ=700 stdev | i>1 max score μ=700 | i>1 max score μ=700 stdev |
|---|---|---|---|---|---|---|---|---|---|---|---|---|---|
| 1 | EPHA5 | 0.002785 | 0.000544 | 0.000893 | 0.00054 | 0.953678 | 0.000488 | 0.956462 | 0.000974 | 0.920503 | 0.000402 | 0.921047 | 0.000742 |
| 2 | BCR | 0.002973 | 0.000188 | 0.000302 | 8e-05 | 0.994297 | 0.000345 | 0.997269 | 0.000136 | 0.972387 | 0.000214 | 0.972575 | 0.000257 |
| 3 | CDK17 | 0.000724 | 0.001855 | 6.5e-05 | 0.00046 | 0.998674 | 4.2e-05 | 0.999398 | 5.1e-05 | 0.983101 | 0.000312 | 0.984956 | 0.00045 |
| 4 | TYK2 | 0.00083 | 0.001339 | 0.000251 | 0.000377 | 0.959997 | 0.000245 | 0.960827 | 0.000206 | 0.93095 | 0.000269 | 0.932288 | 0.000469 |
| 5 | PDPK1;PDPK2P | 0.001601 | -8e-06 | 0.000312 | 0.000341 | 0.960947 | 0.000597 | 0.962548 | 0.000312 | 0.958126 | 0.00033 | 0.958117 | 0.000155 |
| 6 | PRKCD | 0.001281 | 9.4e-05 | 0.000365 | 0.000147 | 0.983142 | 0.000286 | 0.984424 | 0.00018 | 0.975658 | 0.000194 | 0.975752 | 0.00015 |
| 7 | PRKD2 | 0.001184 | 0.00017 | 0.000401 | 0.000433 | 0.963003 | 0.000783 | 0.964186 | 0.000431 | 0.934061 | 0.000297 | 0.934231 | 0.000591 |

**S16. Kinases with significant improvements in off-target effects using combinations of inhibitors from Klaeger *et al* (2017): figure 4 extended data**

Kinase targets were assessed using combinations of inhibitors from Klaeger *et al* (2017). Tight penalty distribution and medium penalty distribution priors were used (see main text Fig 4). Five technical replicates were performed for all analyses with significance for each penalty distribution prior assessed by $p < 0.05$, and an absolute score change > 0.001. Noise distribution variance was set at 2.5, and high on-target penalty set at 0.1.

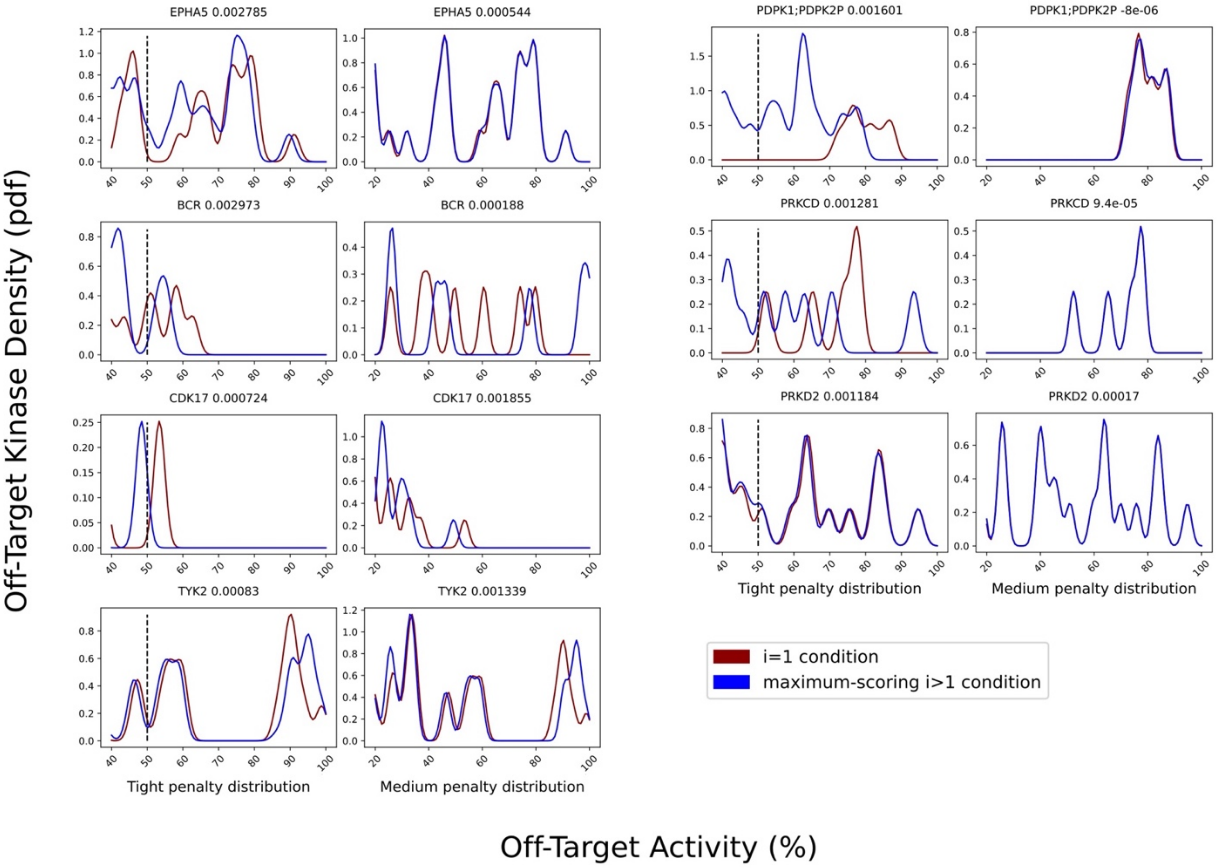


**S17. Off-target activity profiles suggest improvements in off-target effects using inhibitor combinations with inhibitors from Klaeger *et al* (2017): figure 4 extended data**

Probability density functions illustrate the average off-target effects for the highest scoring single inhibitor versus the highest scoring combination of inhibitors for inhibitor combinations from the Klaeger *et al* (2017) dataset, scored using either a tight or medium penalty distribution with an absolute JSD score increase of >=0.001. The variance of the normal distributions used to generate the pdfs is the same as was used in the scoring method: 2.5, such that the overall density is an accurate reflection of the off-target penalty distributions. Vertical dotted lines represent the approximate activity value where the tight probability distribution becomes negligible. Like the tight penalty distribution, the medium penalty distribution penalizes higher off-target effects much more than lower off-target effects, and becomes negligible at approximately 30% activity. Although off-target effects for a kinase may have only been significantly improved using one of the two penalty distributions (tight or medium), data from both is presented to illustrate corresponding non-significant results. Scoring of TYK2 likely represents a false-positive result.

A

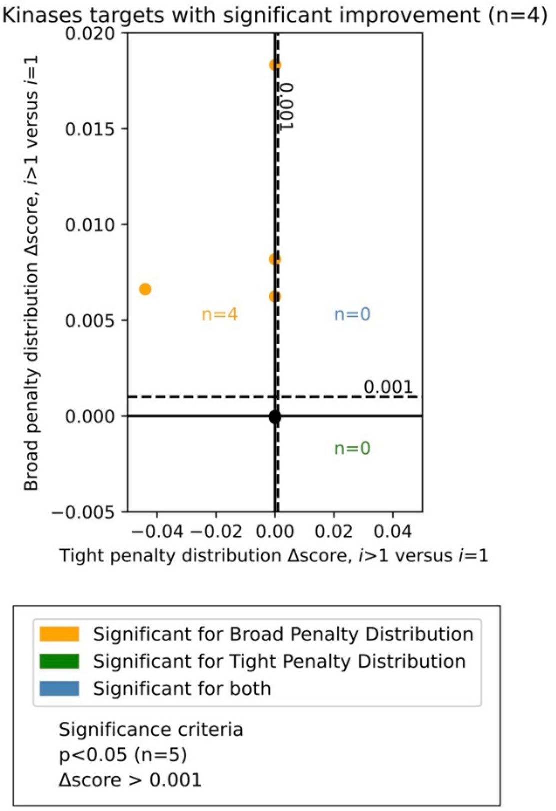


B

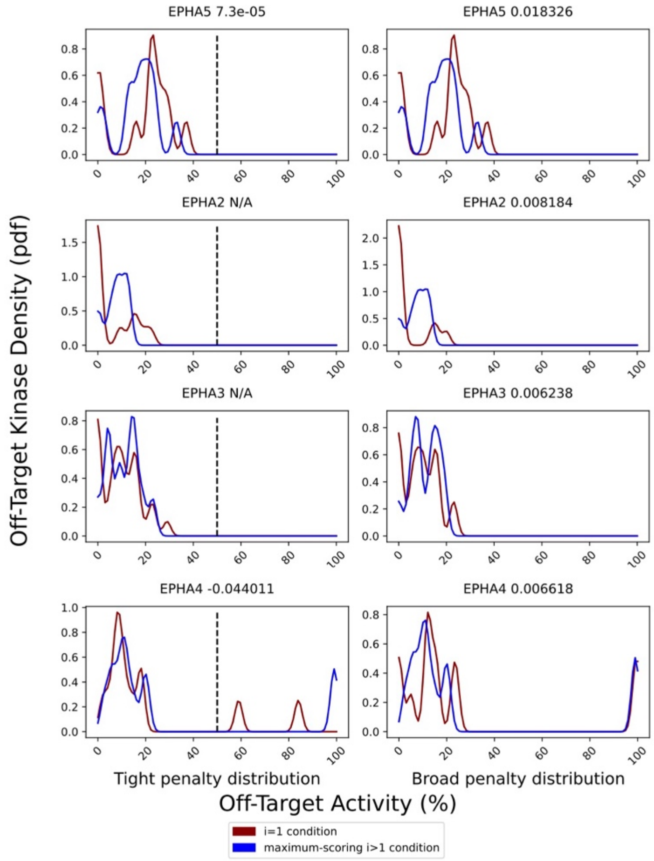


C

| Rank | Kinase | ΔJSD μ=200 | ΔJSD μ=1200 | ΔJSD μ=200 stdev | ΔJSD μ=1200 stdev | i=1 score μ=200 | i=1 score μ=200 stdev | i>1 max score μ=200 | i>1 max score μ=200 stdev |
|---|---|---|---|---|---|---|---|---|---|
| 1 | EPHA5 | 7.3e-05 | 0.018326 | 2.9e-05 | 0.000908 | 0.999891 | 4.9e-05 | 0.999964 | 5.1e-05 |
| 2 | EPHA2 | N/A | 0.008184 | 0.0 | 0.000941 | 1.0 | 0.0 | 1.0 | 0.0 |
| 3 | EPHA3 | N/A | 0.006238 | 0.0 | 0.000883 | 1.0 | 0.0 | 1.0 | 0.0 |
| 4 | EPHB2 | N/A | 3e-06 | 0.0 | 0.000894 | 1.0 | 0.0 | 1.0 | 0.0 |
| 5 | EPHB6 | N/A | -9.6e-05 | 0.0 | 0.000498 | 1.0 | 0.0 | 1.0 | 0.0 |
| 6 | EPHB1 | -8.9e-05 | -0.014178 | 2.6e-05 | 0.000558 | 0.999977 | 2.9e-05 | 0.999888 | 4.9e-05 |
| 7 | EPHA1 | -1e-06 | -0.029931 | 3e-06 | 0.001169 | 0.998908 | 0.000225 | 0.998907 | 0.000226 |
| 8 | EPHA4 | -0.044011 | 0.006618 | 0.004588 | 0.000946 | 0.909126 | 0.002344 | 0.865115 | 0.002315 |
| 9 | EPHA8 | -0.033343 | -0.051397 | 0.002064 | 0.007371 | 0.942921 | 0.002235 | 0.909577 | 0.004092 |
| 10 | EPHA6 | -0.084295 | -0.07914 | 0.001695 | 0.001376 | 0.867848 | 0.002534 | 0.783553 | 0.001322 |
| 11 | EPHB3 | -0.268937 | -0.117277 | 0.006861 | 0.003793 | 0.867848 | 0.002534 | 0.598911 | 0.00902 |
| 12 | EPHA7 | -0.190442 | -0.270131 | 0.001747 | 0.008735 | 0.918963 | 0.001134 | 0.728521 | 0.00259 |
| 13 | EPHB4 | -0.32878 | -0.145505 | 0.010249 | 0.006715 | 0.953509 | 0.001794 | 0.62473 | 0.008717 |

| Rank | Kinase | i=1 score μ=1200 | i=1 score μ=1200 stdev | i>1 max score μ=1200 | i>1 max score μ=1200 stdev |
|---|---|---|---|---|---|
| 1 | EPHA5 | 0.901594 | 0.001836 | 0.91992 | 0.001136 |
| 2 | EPHA2 | 0.961302 | 0.00153 | 0.969486 | 0.000698 |
| 3 | EPHA3 | 0.950944 | 0.000888 | 0.957182 | 0.001416 |
| 4 | EPHB2 | 0.968121 | 0.001362 | 0.968123 | 0.001011 |
| 5 | EPHB6 | 0.978857 | 0.000902 | 0.978761 | 0.000717 |
| 6 | EPHB1 | 0.953629 | 0.001194 | 0.939451 | 0.001382 |
| 7 | EPHA1 | 0.894681 | 0.002474 | 0.86475 | 0.002517 |
| 8 | EPHA4 | 0.839612 | 0.001551 | 0.84623 | 0.001527 |
| 9 | EPHA8 | 0.787471 | 0.005825 | 0.736074 | 0.004846 |
| 10 | EPHA6 | 0.839612 | 0.001551 | 0.760473 | 0.001226 |
| 11 | EPHB3 | 0.839612 | 0.001551 | 0.722336 | 0.004637 |
| 12 | EPHA7 | 0.882774 | 0.003139 | 0.612643 | 0.00848 |
| 13 | EPHB4 | 0.879776 | 0.003154 | 0.734272 | 0.007274 |

**S18. Minor reduction of off-target effects for the EPH kinase family user case**

The PKIS2 dataset was reduced to only the EPH kinase family. Analysis of combinations of up to three inhibitors were performed using either a tight (μ=200, as previously described) or broad (μ=1200) penalty distribution prior, with five technical replicates, and significance assessed by $p < 0.05$, and an absolute score change > 0.001. A JSD score change of N/A is indicated in cases where the single most selective inhibitor had no off-target effects within the range of the penalty distribution, the score was 1, so no further improvements could be made. The variance of the normal distributions used to generate the pdfs is the same as was used in the scoring method: 2.5, such that the overall density is an accurate reflection of the off-target penalty distributions. Vertical dotted lines represent the approximate activity value where the tight probability distribution becomes negligible. Like the tight penalty distribution, the medium penalty distribution penalizes higher off-target effects much more than lower off-target effects, and becomes negligible at approximately 30% activity. Although off-target effects for a kinase may have only been significantly improved using one of the two penalty distributions (tight or medium), data from both is presented to illustrate corresponding non-significant results.

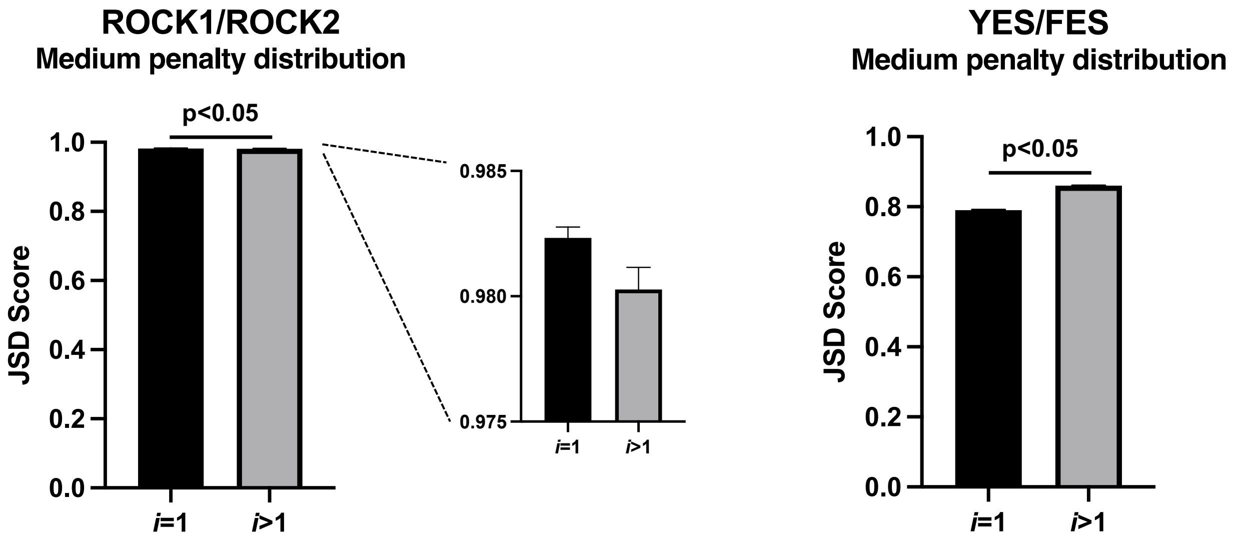


**S19. Fig 6 extended data**

A single inhibitor outperforms combinations of inhibitors for the closely related ROCK1/ROCK2 kinases. A combination of inhibitors outperforms the single most selective inhibitor for the divergent YES/FES kinases. Data plotted as mean +/- SEM for five technical replicates. Analysis performed using a minimum on-target activity threshold for all target kinases of 90%, a noise distribution variance of 2.5, and high off-target penalty of 0.1.

A

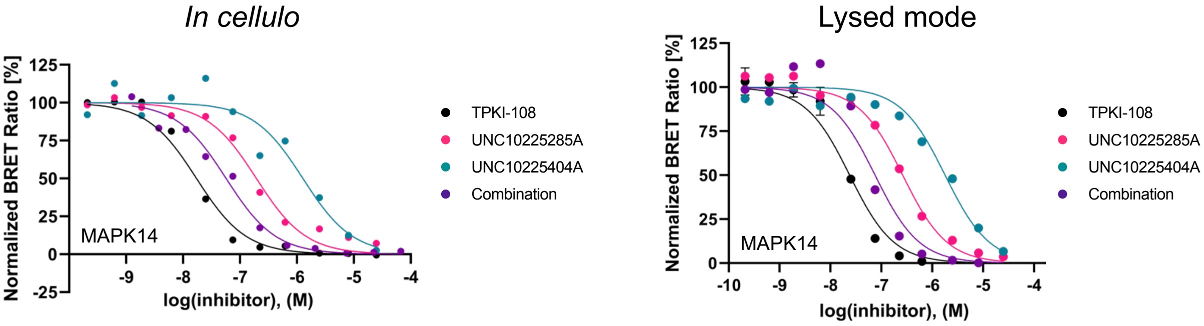


B

**TPKI-108**

| | Rep1 | Rep2 | Rep3 | Rep4 | Rep5 | Average | SD | SEM |
|---|---|---|---|---|---|---|---|---|
| MAPK14 | 1.73E-08 | 1.20E-07 | 1.80E-07 | 3.89E-08 | 5.28E-08 | 8.17E-08 | 5.99E-08 | 2.68E-08 |
| PKN1 | 4.00E-05 | >50 µM | >50 µM | N/A | N/A | 4.00E-05 | N/A | N/A |
| RPS6KA1 | >50 µM | N/A | N/A | N/A | N/A | >50 µM | N/A | N/A |
| RPS6KA6 | 4.44E-05 | >50 µM | >50 µM | >50 µM | >50 µM | 4.44E-05 | N/A | N/A |
| MAPK11 | 8.00E-08 | 6.03E-08 | 9.35E-08 | 1.57E-07 | 2.20E-07 | 1.22E-07 | 5.87E-08 | 2.62E-08 |
| ABL1 | >50 µM | N/A | N/A | N/A | N/A | >50 µM | N/A | N/A |
| BRAF | >50 µM | N/A | N/A | N/A | N/A | >50 µM | N/A | N/A |
| CAMK1 | >50 µM | 4.35E-05 | 4.22E-05 | >50 µM | 4.48E-05 | 4.35E-05 | 1.27E-06 | 7.30E-07 |
| JAK1 | >50 µM | >50 µM | >50 µM | N/A | >50 µM | >50 µM | N/A | N/A |
| KIT | >50 µM | N/A | N/A | N/A | N/A | >50 µM | N/A | N/A |
| LRRK2 | >50 µM | >50 µM | >50 µM | N/A | >50 µM | >50 µM | N/A | N/A |
| PRKCQ | >50 µM | N/A | N/A | N/A | N/A | >50 µM | N/A | N/A |

**UNC10225285A**

| | Rep1 | Rep2 | Rep3 | Rep4 | Rep5 | Average | SD | SEM |
|---|---|---|---|---|---|---|---|---|
| MAPK14 | 1.95E-07 | 1.37E-06 | 6.68E-06 | 3.16E-07 | 4.63E-07 | 1.80E-06 | 2.47E-06 | 1.11E-06 |
| PKN1 | 3.58E-05 | >50 µM | >50 µM | N/A | >50 µM | 3.58E-05 | N/A | N/A |
| RPS6KA1 | >50 µM | N/A | N/A | N/A | N/A | >50 µM | N/A | N/A |
| RPS6KA6 | 2.30E-05 | >50 µM | 3.96E-05 | 4.98E-05 | 3.23E-05 | 3.62E-05 | 9.81E-06 | 4.91E-06 |
| MAPK11 | 5.35E-07 | 1.42E-06 | 5.35E-07 | 1.71E-06 | 1.73E-06 | 1.19E-06 | 5.43E-07 | 2.43E-07 |
| ABL1 | >50 µM | N/A | N/A | N/A | N/A | >50 µM | N/A | N/A |
| BRAF | >50 µM | N/A | N/A | N/A | N/A | >50 µM | N/A | N/A |
| CAMK1 | >50 µM | >50 µM | 3.24E-05 | 4.71E-05 | 3.55E-05 | 3.83E-05 | 6.36E-06 | 3.67E-06 |
| JAK1 | 2.12E-05 | >50 µM | 2.10E-05 | 3.69E-05 | 3.59E-05 | 2.87E-05 | 7.66E-06 | 3.83E-06 |
| KIT | >50 µM | N/A | N/A | N/A | N/A | >50 µM | N/A | N/A |
| LRRK2 | 7.27E-06 | >50 µM | >50 µM | N/A | 8.48E-06 | 7.87E-06 | 6.06E-07 | 4.29E-07 |
| PRKCQ | >50 µM | N/A | N/A | N/A | N/A | >50 µM | N/A | N/A |

**UNC10225404A**

| | Rep1 | Rep2 | Rep3 | Rep4 | Rep5 | Average | SD | SEM |
|---|---|---|---|---|---|---|---|---|
| MAPK14 | 1.28E-06 | 8.90E-06 | 9.25E-06 | 1.97E-06 | 1.73E-06 | 4.62E-06 | 4.07E-06 | 1.82E-06 |
| PKN1 | >50 µM | >50 µM | >50 µM | N/A | N/A | >50 µM | N/A | N/A |
| RPS6KA1 | >50 µM | N/A | N/A | N/A | N/A | >50 µM | N/A | N/A |
| RPS6KA6 | >50 µM | >50 µM | >50 µM | >50 µM | >50 µM | >50 µM | N/A | N/A |
| MAPK11 | 2.03E-06 | 1.48E-06 | 1.67E-06 | 3.64E-06 | 3.23E-06 | 2.41E-06 | 9.68E-07 | 4.33E-07 |
| ABL1 | >50 µM | N/A | N/A | N/A | N/A | >50 µM | N/A | N/A |
| BRAF | >50 µM | N/A | N/A | N/A | N/A | >50 µM | N/A | N/A |
| CAMK1 | >50 µM | >50 µM | 4.57E-05 | >50 µM | >50 µM | 4.57E-05 | N/A | N/A |
| JAK1 | >50 µM | 4.64E-05 | >50 µM | >50 µM | >50 µM | 4.64E-05 | N/A | N/A |
| KIT | >50 µM | N/A | N/A | N/A | N/A | >50 µM | N/A | N/A |
| LRRK2 | >50 µM | >50 µM | >50 µM | N/A | 4.48E-05 | 4.48E-05 | N/A | N/A |
| PRKCQ | >50 µM | N/A | N/A | N/A | N/A | >50 µM | N/A | N/A |

**Combination**

| | Rep1 | Rep2 | Rep3 | Rep4 | Rep5 | Average | SD | SEM |
|---|---|---|---|---|---|---|---|---|
| MAPK14 | 5.72E-08 | 3.99E-07 | 5.78E-07 | 7.21E-08 | 1.25E-07 | 2.46E-07 | 2.31E-07 | 1.03E-07 |
| PKN1 | >50 µM | >50 µM | >50 µM | N/A | N/A | >50 µM | N/A | N/A |
| RPS6KA1 | >50 µM | N/A | N/A | N/A | N/A | >50 µM | N/A | N/A |
| RPS6KA6 | 3.98E-05 | >50 µM | >50 µM | >50 µM | >50 µM | 3.98E-05 | N/A | N/A |
| MAPK11 | 2.87E-07 | 3.42E-07 | >50 µM | 6.57E-07 | 5.86E-07 | 4.68E-07 | 1.81E-07 | 9.05E-08 |
| ABL1 | >50 µM | N/A | N/A | N/A | N/A | >50 µM | N/A | N/A |
| BRAF | >50 µM | N/A | N/A | N/A | N/A | >50 µM | N/A | N/A |
| CAMK1 | 8.73E-06 | >50 µM | 2.66E-05 | 4.17E-05 | 3.31E-05 | 2.75E-05 | 1.40E-05 | 6.98E-06 |
| JAK1 | >50 µM | >50 µM | >50 µM | N/A | >50 µM | >50 µM | N/A | N/A |
| KIT | >50 µM | N/A | N/A | N/A | N/A | >50 µM | N/A | N/A |
| LRRK2 | >50 µM | >50 µM | >50 µM | >50 µM | >50 µM | >50 µM | N/A | N/A |
| PRKCQ | >50 µM | N/A | N/A | N/A | N/A | >50 µM | N/A | N/A |

C

**Replicate 1 lysed**

| | TPKI-108 | UNC10225285A | UNC10225404A | Combination |
|---|---|---|---|---|
| LRRK2 | N/A | N/A | N/A | N/A |
| MAPK14 | 4.24E-08 | 5.99E-07 | 2.45E-06 | 1.34E-07 |
| JAK1 | 3.83E-05 | 5.17E-05 | 7.39E-05 | 6.51E-05 |
| CAMK1 | 3.07E-05 | 8.48E-05 | 1.25E-04 | 5.70E-05 |
| PKN1 | 7.26E-05 | Unstable | Unstable | 3.20E-04 |
| RPS6KA6 | 5.34E-05 | 6.00E-04 | 1.19E-04 | 3.86E-04 |
| MAPK11 | 1.04E-07 | 8.38E-07 | 4.05E-06 | 4.12E-07 |

**Replicate 2 lysed**

| | TPKI-108 | UNC10225285A | UNC10225404A | Combination |
|---|---|---|---|---|
| LRRK2 | N/A | N/A | N/A | N/A |
| MAPK14 | 3.14E-08 | 2.63E-07 | 1.79E-06 | 8.96E-08 |
| JAK1 | 8.49E-05 | 9.23E-05 | 1.08E-04 | 7.60E-05 |
| CAMK1 | 2.61E-05 | 4.50E-05 | 9.12E-05 | 2.87E-05 |
| PKN1 | 2.06E-05 | 2.27E-05 | 9.49E-04 | 2.18E-05 |
| RPS6KA6 | 1.52E-04 | 4.51E-04 | 5.08E-02 | 3.53E-04 |
| MAPK11 | 9.03E-08 | 5.66E-07 | 3.41E-06 | 3.02E-07 |

**Average lysed**

| | TPKI-108 | UNC10225285A | UNC10225404A | Combination |
|---|---|---|---|---|
| LRRK2 | N/A | N/A | N/A | N/A |
| MAPK14 | 3.69E-08 | 4.31E-07 | 2.12E-06 | 1.12E-07 |
| JAK1 | 6.16E-05 | 7.20E-05 | 9.11E-05 | 7.05E-05 |
| CAMK1 | 2.84E-05 | 6.49E-05 | 1.08E-04 | 4.29E-05 |
| PKN1 | 4.66E-05 | 2.27E-05 | 9.49E-04 | 1.71E-04 |
| RPS6KA6 | 1.02E-04 | 5.25E-04 | 2.54E-02 | 3.69E-04 |
| MAPK11 | 9.73E-08 | 7.02E-07 | 3.73E-06 | 3.57E-07 |

**SD lysed**

| | TPKI-108 | UNC10225285A | UNC10225404A | Combination |
|---|---|---|---|---|
| LRRK2 | N/A | N/A | N/A | N/A |
| MAPK14 | 5.49E-09 | 1.68E-07 | 3.30E-07 | 2.20E-08 |
| JAK1 | 2.33E-05 | 2.03E-05 | 1.73E-05 | 5.42E-06 |
| CAMK1 | 2.27E-06 | 1.99E-05 | 1.69E-05 | 1.42E-05 |
| PKN1 | 2.60E-05 | N/A | N/A | 1.49E-04 |
| RPS6KA6 | 4.91E-05 | 7.48E-05 | 2.53E-02 | 1.63E-05 |
| MAPK11 | 6.99E-09 | 1.36E-07 | 3.21E-07 | 5.52E-08 |

**S20. NanoBRET *in cellulo* and lysed mode compound and combination activity: figure 5 extended data**

The cumulative activity each compound and the equimolar combination of compounds was studied via an 11-point in-cell NanoBRET target engagement assay for MAPK14 and the set of major of-targets defined by the PKIS2 MMS analysis. The assay was performed *in cellulo* (A, B) and using cell lysate (A, C). A conservative 50 µM cutoff was imposed for reporting all *in cellulo* $EC_{50}$ values. A representative trace for MAPK14 from both experiments is indicated (A).

A

$$I^{T} = \frac{\sum_{j=1}^{n} \frac{I_j}{K_{ij}}}{1 + \sum_{j=1}^{n} \frac{I_j}{K_{ij}}}$$

$$I^{T} \times 100\% = Activity$$

$$\left(\left(\frac{1}{I^{T}}\right) - 1\right)\sum_{j=1}^{n} I_j = EC_{50}(M)$$

B

| *in cellulo* | Combination Activity at 1 µM (calculated) | Combination $EC_{50}$ (calculated) | Combination $EC_{50}$ (observed) |
|---|---|---|---|
| **MAPK14** | 81.25 | 2.31E-07 | 2.46E-07 |
| **PKN1** | N/A | N/A | >50 µM |
| **RPS6KA6** | N/A | N/A | 3.98E-05 |
| **MAPK11** | 75.89 | 3.18E-07 | 4.68E-07 |
| **CAMK1** | 2.31 | 4.23E-05 | 2.75E-05 |
| **JAK1** | N/A | N/A | >50 µM |
| **LRRK2** | N/A | N/A | >50 µM |

C

| Lysed mode | Combination Activity at 1 µM (calculated) | Combination $EC_{50}$ (calculated) | Combination $EC_{50}$ (observed) |
|---|---|---|---|
| **MAPK14** | 90.88 | 1.00E-07 | 1.12E-07 |
| **PKN1** | 2.17 | 4.51E-05 | 1.71E-04 |
| **RPS6KA6** | 0.39 | 2.56E-04 | 3.69E-04 |
| **MAPK11** | 79.96 | 2.51E-07 | 3.57E-07 |
| **CAMK1** | 1.96 | 5.01E-05 | 4.29E-05 |
| **JAK1** | 1.35 | 7.30E-05 | 7.05E-05 |
| **PKN1** | 2.17 | 4.51E-05 | 1.71E-04 |

**S21. Calculation of cumulative NanoBRET *in cellulo* and lysed mode combination activity**

The cumulative activity of the three compounds was calculate for each kinase using the same method as in the MMS protocol (see supplementary methods) and using the average $EC_{50}$ of each compound for each kinase. The cumulative activity and $EC_{50}$ is given by the equations in (A) where $I^T$ is the total inhibition of a kinase by n inhibitors, $I_j$ is the concentration of an inhibitor *j*, and $K_{ij}$ is the $K_i$ or $K_i$ equivalent of inhibitor *j* against the target kinase. Percentages are used for the activity scale. The cumulative activity and cumulative $EC_{50}$s for both the *in cellulo* (B) and lysed mode (C) experiments are indicated, along with the average observed $EC_{50}$s. N/A indicates that a value could not be calculated due to $EC_{50}$s > 50 µM for one or more of the single inhibitors.

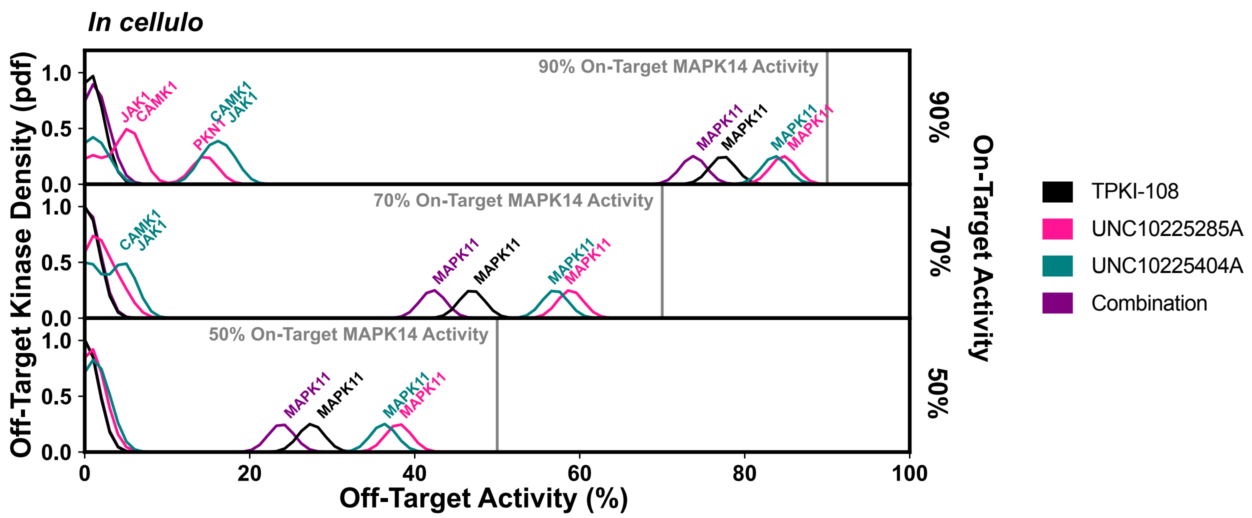


**S22. Validation of compound combination predicted *in cellulo* selectivity: figure 5 extended data**

Calculation of the off-target profiles of the inhibitor combination and the individual inhibitors from *in cellulo* $EC_{50}$s suggests that the combination modestly improves selectivity for MAPK14 over the most selective single inhibitor, TPKI-108. The off-target activity of the inhibitors is calculated based upon their $EC_{50}$s for each off-target kinase, at the concentration needed to reach 90%, 70% or 50% on-target activity against MAPK14. The variance of the normal distributions used to generate probability density functions (pdfs) is the same as was used in the MMS scoring method (2.5) and the activity scale is shown between 0 and 100.

**Table 1. Kinase co-targets more selectively inhibited by a combination of more than two inhibitors than by either the single most selective inhibitor or the most selective combination of two inhibitors**

ΔJSD score: max($i$=3,$i$=4) - max($i$=1,$i$=2). Kinases were selected for analysis from a single-target screen of the Klaeger et al dataset if they had a single-compound JSD score <0.95. All possible pairs of kinases from this set were scored, up to $i$=4. Pairs that were significant according to MMS protocol (absolute cutoff=0.001) with both penalty prior shapes were curated. The sum score is the total JSD score using both priors.

| | **Kinase Targets** | **ΔJSD score tight prior** | **ΔJSD score medium prior** | **sum score** |
|---|---|---|---|---|
| 1 | FER_PLK4 | 0.04275 | 0.03710 | 0.07985 |
| 2 | PRKD2_CSNK1G1 | 0.02804 | 0.04276 | 0.07080 |
| 3 | FER_PTK2B | 0.03017 | 0.03369 | 0.06386 |
| 4 | TBK1_STK3 | 0.02574 | 0.03613 | 0.06187 |
| 5 | TANK_STK3 | 0.02628 | 0.03433 | 0.06062 |
| 6 | PTK6_FER | 0.02862 | 0.03069 | 0.05931 |

| | | | | |
|---|---|---|---|---|
| 7 | BUB1_TGFBR2 | 0.02579 | 0.02767 | 0.05347 |
| 8 | PRKD3_MAPK15 | 0.02343 | 0.02692 | 0.05035 |
| 9 | RPS6KA4_TP53RK | 0.02111 | 0.02733 | 0.04845 |
| 10 | LYN_PTK6 | 0.01691 | 0.02501 | 0.04192 |
| 11 | PLK4_TGFBR2 | 0.01735 | 0.01032 | 0.02768 |
| 12 | STK3_MAPK15 | 0.00942 | 0.01821 | 0.02764 |
| 13 | CDK4_PRKD2 | 0.01278 | 0.00727 | 0.02005 |
| 14 | RPS6KA3_TBK1 | 0.00943 | 0.00998 | 0.01941 |
| 15 | STK4_MAP4K1 | 0.00253 | 0.01634 | 0.01887 |
| 16 | TAOK3_MELK | 0.00830 | 0.00953 | 0.01783 |
| 17 | RPS6KA4_TPRKB | 0.00627 | 0.01020 | 0.01647 |
| 18 | PLK4_ACVRL1 | 0.00905 | 0.00661 | 0.01567 |
| 19 | CDK1_CDK7 | 0.00933 | 0.00584 | 0.01517 |
| 20 | LCK_CSK | 0.00483 | 0.01015 | 0.01498 |
| 21 | PTK6_EPHB6 | 0.00626 | 0.00711 | 0.01337 |
| 22 | MAP3K11_ULK1 | 0.00738 | 0.00585 | 0.01323 |
| 23 | ERN1_STK3 | 0.00720 | 0.00539 | 0.01259 |
| 24 | RPS6KA4_PRKG1 | 0.00389 | 0.00845 | 0.01234 |
| 25 | PRKD3_ERN1 | 0.00547 | 0.00668 | 0.01214 |
| 26 | TAOK3_PTK2B | 0.00670 | 0.00478 | 0.01149 |
| 27 | FER_PIP4K2C | 0.00468 | 0.00677 | 0.01145 |

| | | | | |
|---|---|---|---|---|
| 28 | MAP3K11_NUAK2 | 0.00514 | 0.00630 | 0.01144 |
| 29 | TNIK_MARK4 | 0.00374 | 0.00754 | 0.01127 |
| 30 | STK10_LYN | 0.00585 | 0.00497 | 0.01082 |
| 31 | PRKD2_ERN1 | 0.00559 | 0.00492 | 0.01050 |
| 32 | EPHA4_PLK4 | 0.00441 | 0.00574 | 0.01015 |
| 33 | RPS6KA6_EPHA7 | 0.00276 | 0.00677 | 0.00952 |
| 34 | PRKD2_MELK | 0.00565 | 0.00357 | 0.00922 |
| 35 | MAP4K4_ULK1 | 0.00132 | 0.00783 | 0.00914 |
| 36 | STK4_BUB1 | 0.00599 | 0.00296 | 0.00895 |
| 37 | ULK3_MAP2K6 | 0.00318 | 0.00517 | 0.00834 |
| 38 | MAP4K4_STK3 | 0.00184 | 0.00568 | 0.00752 |
| 39 | CAMK2G_TBKBP1 | 0.00251 | 0.00491 | 0.00742 |
| 40 | TNIK_STK3 | 0.00302 | 0.00437 | 0.00739 |
| 41 | EPHB3_TGFBR2 | 0.00444 | 0.00271 | 0.00715 |
| 42 | FRK_PLK4 | 0.00165 | 0.00544 | 0.00709 |
| 43 | TNIK_CLK4 | 0.00423 | 0.00279 | 0.00702 |
| 44 | PRKAA1_MARK4 | 0.00444 | 0.00216 | 0.00660 |
| 45 | SIK2_ACVRL1 | 0.00156 | 0.00498 | 0.00654 |
| 46 | PAK4_MARK3 | 0.00266 | 0.00382 | 0.00649 |
| 47 | SIK2_STK4 | 0.00348 | 0.00242 | 0.00590 |
| 48 | ZAK_LYN | 0.00108 | 0.00431 | 0.00539 |

| | | | | |
|---|---|---|---|---|
| 49 | MYLK_NLK | 0.00330 | 0.00199 | 0.00530 |
| 50 | SIK2_STK3 | 0.00164 | 0.00359 | 0.00523 |
| 51 | CLK4_IRAK4 | 0.00282 | 0.00236 | 0.00518 |
| 52 | CIT_TESK1 | 0.00308 | 0.00167 | 0.00474 |
| 53 | LCK_FGR | 0.00265 | 0.00159 | 0.00424 |
| 54 | PRKAG2_MARK4 | 0.00222 | 0.00114 | 0.00336 |
| 55 | EPHB3_LCK | 0.00101 | 0.00114 | 0.00215 |
| 56 | EPHB4_TESK1 | 0.00109 | 0.00103 | 0.00213 |

**Supplementary Methods**

*Multicompound-multitarget scoring (MMS) method*

First, either a single kinase, or a group of multiple kinases, are defined as the target set. The inhibitors with potent activity against these targets, over a user-defined threshold, are divided into combinations of $i$ inhibitors, where $i$ is the number of inhibitors in a single combination. For each combination number $i$, there are $c\_i$ unique combinations of inhibitors. If the target set contains multiple kinases, combinations that do not maintain 90% activity for all target kinases are eliminated. We use an inhibitor activity threshold of 90% for all experiments, which corresponds with a single-inhibitor $K_i$ of at least 111 nM if compounds are dosed in a reference frame of 1 µM. Program settings also facilitate input of inhibitor $K_i$ values, $K_d$ values, or $K_d^{app}$ values, which are treated equivalently.

Next, the user defines a selectivity penalty distribution. This is a probability distribution that differentially penalizes off-target effects, with a maximum at 100% activity and a left-skewed tail. Users can control the curvature of this distribution and where the tail ends, thus defining the gradient of weights that should be applied as well as the minimum off-target effect relevant to their goals. For example, if compounds with off-target effects less than 50% activity are not meaningful in a user-specific context, that user may opt to use a tight penalty distribution with minimal left-skew that is relatively insensitive to off-target effects below that threshold. Alternatively, if all off-target effects are meaningful to a user, they may opt to use a broad penalty prior that is highly left-skewed, which includes some penalty for even very low off-target inhibitor activities. We compare distributions based upon common distributions shapes, either the 1) beta probability density function or 2) the left-tail of the poisson probability mass function, given by:

1. $\frac{\Gamma(\alpha+\beta)x^{\alpha-1}(1-x)^{\beta-1}}{\Gamma(\alpha)\Gamma(\beta)}$ for $0 \leq x \leq 1$ where Γ( ) is the gamma function, for shape parameters $\alpha$ and $\beta$ where $\alpha$>0 and $\beta$>0
2. $\frac{\mu^k e^{-\mu}}{k!}$ for $k \geq \mathbf{0}$, for shape parameter $\mu$ where $\mu \geq 0$

We find that, for equivalent scores generated by the distributions, there is only a minimal difference in the performance of the penalty priors (Fig S1; S2). For the distributions based upon the poisson prior there is no correlation between results reproducibility and score, versus a minor correlation for the distributions based upon beta priors (Fig S2). For this reason, the poisson prior shape is selected as the basis for all following analyses, altering the skew of the distribution to produce either a tight, medium, or broad penalty prior. The center of the poisson prior (controlled by the μ value) is placed at the equivalent position to 100% activity, or 1 in the normalized reference frame, and only the left tail of the distribution is used. See main text Fig 2 for a visualization. Therefore, in practice the μ value controls the overall spread of the poison prior. These penalty priors all penalize off-target effects closer to 100% activity more than the lowest off-target effects recognized by the distributions, although the tight penalty prior has the greatest penalties for high off-target activity. Similarly, the broad penalty distribution has the lowest penalties for high off-target activity. We also integrate the option to include an additional penalty to the highest set of off-target effects, defined for these analyses as the 95%-100% activity range. This high off-target penalty accounts for logarithmic fold-changes in $K_i$, $K_d$, or $K_d^{app}$ that are not readily apparent in the linear activity-threshold scale, and does not affect method performance (Fig S1). We conduct complementary analyses using two sets of penalty priors for all datasets: a tight penalty prior and either a medium or broad penalty prior, and recommend that similar complementary analyses should be performed for unique cases of datasets and targets. A single prior could be used in user cases where a particular range of off-target effects is of particular interest; however, we suggest using complementary priors for general analyses if such information is not known or accessible. Inhibitor combinations may only be significant with one of two complementary prior shapes; in these cases it is important to consider context, interpretation, and user goals.

Once all inhibitor combinations and the penalty distribution have been generated, the cumulative activity of all inhibitors in a combination *c_i* against each off-target kinase is calculated. The cumulative activity of multiple competitive compounds at a single target is given by:

3. $\mathrm{I}^{\mathrm{T}} = \frac{\sum_{j=1}^{n} \frac{I_j}{K_{ij}}}{1 + \sum_{j=1}^{n} \frac{I_j}{K_{ij}}}$ where $I^T$ is the total inhibition of a kinase by n inhibitors, $I_j$ is the concentration of an inhibitor *j*, and $K_{ij}$ is the $K_i$ or $K_i$ equivalent of inhibitor *j* against the target kinase. Percentages are used for the activity scale. We note that this is an approximation that does not include the effect of varying the concentration of ATP or the kinase. See Figure S21 for the form used to generate the equivalent combination $K_i$ (or NanoBRET $EC_{50}$).

Each single cumulative activity calculation is used as the mean for a Gaussian distribution from which noise is bootstrapped. The variance of this noise distribution can be defined by the user. Increasing the variance of this distribution decreases the reproducibility of program results (Fig S1), but accounts for uncertainty in data measurements. Program reproducibility is defined as the percentage of first-rank identical inhibitor sets obtained from a set of five technical replicates – variability in results is introduced by increasing the variance of the noise distribution, in addition to resampling the penalty distribution between technical replicates. Consequently, program reproducibility decreases with increasing values of *i* to approximately 60% for combinations of *i*=3 inhibitors (Fig S1). Non-perfect reproducibility is an indication of uncertainty in the results, and suggests that other equally selective single inhibitors or inhibitor combinations may be available. Reproducibility is not a measure of method validity.

Off-target effects are combined into a single off-target distribution. This off-target distribution and the penalty distribution are both normalized and binned by off-target inhibitor activity ranges. We use a bin size of 5% activity as an analytical compromise; too small a bin size would make scoring overly sensitive to variations in noise sampling, and too large a range would be insensitive to functionally meaningful differences in off-target profiles. Additionally, intervals of 5% are commonly used for kinase activity profiles [24]. While inhibitor activity range units are used in this implementation to describe the functional relevance of inhibitor action, it is also possible to integrate scales based on $K_i$, $K_d$, or $K_d^{app}$ values.

The Jensen-Shannon distance (JSD score) between the penalty distribution and the off-target distribution is calculated. To summarize, the penalty distribution is a normalized probability distribution with a shape defined by the user. The off-target distribution describes the set of off-target kinases, where each kinase is represented by the cumulative activity of the set of inhibitor(s) in use against that kinase (on our 0-100% activity scale), and each cumulative activity calculation is used as the mean value for a Gaussian distribution from which values are bootstrapped, and those noise values are binned and normalized. The JSD score describes the overlap between these two normalized distributions: the penalty distribution (sometimes referred to as penalty prior) and the off-target distribution. This metric falls between 0 and 1, describing either identical or entirely non-overlapping distributions, respectively. A high score, the product of minimal overlap between the off-target distribution and the penalty distribution, represents a selective inhibitor, or combination of inhibitors. Tight penalty priors tend to produce higher scores than medium or broad priors for equivalent off-target distributions, since fewer off-target effects are included in the calculations (Fig S1). In some cases, there may be no off-target effects in the range that a particular penalty prior is sensitive to, yielding a score of 1, in which case using a broader penalty prior is indicated.

The concentration of inhibitors in each combination $c_i$ is then optimized. Cutoffs can be used to limit negligibly small concentrations of inhibitors, and while we implement no maximum concentration cutoff to account for compound solubility issues we note that such poorly potent compounds would not normally be included in MMS analyses since they would fail to reach 90% on-target activity in a reference frame of 1 µM. Decreasing this cutoff or changing the concentration reference frame could introduce these poorly potent compounds, and is something that a user should consider should they choose to do so. Only the JSD score is maximized during the optimization process, and concentrations that fail to reach on-target threshold activity are excluded. This scoring approach specifically minimizes weighted off-target effects, while retaining on-target activity, unlike relative selectivity measures. Additionally, this process results in dilution of inhibitors down to minimum on-target activity, including for the $i$=1 case, providing a calculation with functional relevance.

Lastly, the performance of the highest-scoring single inhibitor at *i*=1 is compared to the best performing combination for all other values of *i*. The maximum-scoring single inhibitor for the target(s), at *i*=1, represents the most selective single inhibitor. If a combination at *i*>1 has a greater JSD score, or a positive ΔJSD score, at their defined concentrations, then the combination would produce fewer off-target effects than the single most selective inhibitor while retaining potent activity against the target kinase(s). We conduct five technical replicates for each experiment in order to ensure statistical significance, and use an additional absolute ΔJSD score cutoff of 0.001 to eliminate statistically significant but functionally insignificant results. Raising the JSD cutoff to 0.005 correspondingly increases the magnitude of the observed reduction in off-target effects (Fig 3). It is necessary to conduct technical replicates since we introduce noise during the course of the method when we sample points to generate our off-target distribution from normal distributions centered on the calculated activity values of off-target kinases. Additionally, the penalty distribution is also resampled between runs.

*Dataset Preparation*

The Karaman *et al*, Davis *et al*, and Klaeger *et al* datasets of compound $K_d$ or $K_d^{app}$ (nM) were converted to activity values at a reference concentration of 1 µM using the approximation: activity = 100% / [($K_d$/1,000 (M))+1]. A $K_d$ of 100 nM corresponds to an activity approximation of 91%, just above the lower threshold of what is considered a potent compound (90%) in our analyses. The reference frame of 1 µM does not influence calculations and was only used to standardize input. These activity values were converted back into the original $K_d$ or $K_d^{app}$ values by the program during calculations, the changes were only necessary to facilitate data input. PKIS2-activity values were used as direct input for the program.

*Single-Target Analysis*

Single-target analysis was performed as described in the methods section. The following program settings were used: tight prior μ=200, medium prior μ=700, broad prior μ=1200, prior sample size = 100,000, optimization steps R1=5, R2=1.1, on-target threshold = 90, noise distribution variance = 2.5, high off-target penalty = 0.1. Five technical replicates were performed. Significance was assessed by both statistical significance (t-test, $p<0.05$) and an absolute JSD score cutoff. A cutoff of 0.001 was used for all analyses, except when indicated (at 0.005) during simulated data analysis. Kinase mutants in all datasets, while considered as targets for selectivity optimization, were excluded from all calculations of off-target distributions.

*Multiple-target Analysis*

Multiple-target analyses were performed for YES/FES and ROCK1/ROCK2 using the same settings as for single-target analyses. Multiple target analyses were performed for the Klaeger *et al* dataset with the following single settings change: a high off-target penalty of 0.3. A non-exhaustive screen of kinase target pairs from the Klaeger *et al* dataset was performed using the following settings changes: optimization steps R1=3, R2=1.1, and high off-target penalty = 0.3. Kinases with less than a 0.95 single-target JSD score using the medium penalty prior were included in the analysis.

*Eph Family Analysis*

PKIS2 was reduced to only the data for the EPH kinase family, such that only off-target effects for other EPH kinases would be considered for each respective on-target EPH kinase. This data was analyzed using the single-target protocol, with a tight penalty prior (μ=200), and a broad penalty prior (μ=1200), instead of the medium penalty prior (μ=700), in order to capture changes in low off-target effects.

*Program Parameters*

1. Penalty prior type, shape, and size

Both a Poisson distribution and a left-tailed beta distribution can be set by program users, with control over the shape parameters using the μ parameter for the Poisson distribution or $\alpha$ and $\beta$ parameters for the beta distribution. Both distributions are scaled from 0 to 100% activity with an AUC of 1. We observe in our parameter scan tests (Fig S1; S2) that both shapes give reasonably similar reproducibility per score, although there may be slightly better combinatorial resolution when using Poisson distributions for analyses in which a user wants to account more for lower off-target effects. Due to this slight difference, we select the Poisson distribution as the basis of our analyses. We use μ values of 200 or 700 for the majority of our analyses, reflecting the desire to reduce either high-off target effects, or both high and middling off-target effects, respectively. We name these two penalty priors the tight and medium priors in the main text. In one instance we use a μ value of 1200, reflecting the need to weight very low off-target effects. We name this prior the broad prior in the main text. We sample priors using a default of 100,000 points; since the distributions are normalized to an AUC of 1 regardless of their size this number can be increased if desired. We do not recommend making this number too small, as this may result in the penalty prior having odd features, such as non-decreasing step magnitudes with decreasing activity values.

2. High off-target effect penalty

In addition to the shape of the initial penalty distribution, we add an option to increase the penalty of the highest off-target effects, or those at or above 95% compound activity. We find that adding this additional penalty both actualizes the goal of the method, to reduce the highest off-target effects, and helps account for the scaling differences between $K_d$ values and activity approximations or $K_i$ values and activity calculations, at very high $K_d$ or $K_i$ values. For example, a single compound with a $K_i$ of 52nM against an off-target kinase has an estimated activity of 95.05%, while another compound with a $K_i$ of 5.2pM has an estimated activity of 99.99%, and while using a bin size of 5 both would be penalized by the same amount if no variance is introduced to the calculations; see Distribution bin size and Gaussian noise below. This additional penalty is normalized, along with the underlying penalty prior, to an AUC of 1. In multi-target analyses it may be necessary to increase this penalty above 0.1.

3. Distribution bin size

We opt to use a bin size of 5% activity to build our penalty and off-target distributions as a reasonable balance between competing technical limitations and limitations of results interpretation. We note that using a larger bin size would mask changes that a user might find meaningful. Using a smaller bin size would be more responsive to fluctuations in the off-target distribution following the addition of sampled noise measurements, and decrease the reproducibility of results. We find that using a 5% bin size is a good balance between a normative intuition of what constitutes a meaningful change in activity, and a desire to maintain reasonable reproducibility of results following the addition of noise in the program. We also note that, since each measurement is replaced by noise sampled from a normal distribution, the absolute cutoffs of the bins and activity measurements close to these values (ex: 94.8 versus 95.2) do not constitute meaningful differences in the context of program scoring, since sampling will add a similar number of counts to each bin. The bins are primarily useful for smoothing variations in sampling within normative common-sense ranges.

4. Gaussian noise

The default variance for the gaussian distribution from which noise is sampled is 2.5, which complements the program bin size of 5% activity. Increasing the variance decreases the reproducibility of the results generated by the program, or increases the likelihood that alternate equally selective compounds or combinations of compounds will be identified. However, variability is desirable as it reflects uncertainty in input data measurements.

5. On-target inhibition threshold

In all our analyses we maintain an on-target inhibition threshold of 90% activity, representing potent inhibition of target kinases. This threshold controls several points in the program, including how many compounds are initially selected as potent against a target kinase, how far compound combinations are diluted if they have greater than threshold activity, and the minimum activity that must be maintained during compound concentration optimization steps. Decreasing this threshold increases the number of possible compound combinations, and will increase processing time.

6. Compound concentration optimization and optimization step size

We primarily use optimization steps of R1=5 and R2=1.1. We find that an R1 value of 2-5 is useful; higher steps sizes decrease processing time but reduce precision. We do not suggest changing the R2 step size. The step sizes correspond with fold-changes in compound concentration during optimization compound rounds. In the first step of each round, the concentration of each combination of two compounds are varied by R1-fold, leading to a set size of new concentration variations equivalent to the simple choose-two combination. The concentrations of all compounds in these sets are then adjusted by iterative step sizes of R1 to reach minimum on-target activity, usually 90%. Then, the JSD score is calculated, and the top-scoring set of concentrations is used as the input for the next optimization round. Optimization continues until the JSD score can no longer be increased, and the highest-scoring concentrations of the compounds are selected.

*NanoBRET target engagement assay*

The assay was performed as described previously [40]. In brief: Full-length kinase ORF (Promega) cloned in frame with a NanoLuc-vector (as indicated in table below) was transfected into HEK293T cells using FuGENE HD (Promega, E2312) and proteins were allowed to express for 20h. Serially diluted inhibitor and NanoBRET$^{TM}$ Kinase Tracer (as indicated in the table below) were pipetted into white 384-well plates (Greiner 781 207) using an ECHO 550 acoustic dispenser (Labcyte). The corresponding transfected cells were added and reseeded at a density of 2 x $10^5$ cells/ml after trypsinization and resuspension in Opti-MEM without phenol red (Life Technologies). The system was allowed to equilibrate at 37°C for 2h and 5% $CO_2$ prior to BRET measurements. To measure BRET, NanoBRET$^{TM}$ NanoGlo Substrate + Extracellular NanoLuc Inhibitor (Promega, N2160) were added as per the manufacturer's protocol, and filtered luminescence was measured on a PHERAstar plate reader (BMG Labtech) equipped with a luminescence filter pair (450 nm BP filter (donor) and 610 nm LP filter (acceptor)). Competitive displacement data were normalized and then plotted using GraphPad Prism 9 software using a normalized 3-parameter curve fit with the following equation: Y=100/(1+10^((X-LogIC50))). For the PRKCQ assay, the protein was stimulated by adding a final concentration of 1 µM PMA (phorbol 12-myristate 13-acetate, Sigma #P8139) in DMSO.

| Target | Nluc Placement | Target Catalog No | Tracer | [Tracer], [M] | Tracer Catalog No |
|---|---|---|---|---|---|
| MAPK14 | C | NV1661 | K4 | 3.10E-08 | N2540 |
| PKN1 | C | Kind gift of Promega | K16 | 1.30E-07 | Kind gift of Promega |
| RPS6KA1 | N | NV1981 | K10 | 6.30E-08 | N2840 |
| RPS6KA6 | N | NV2021 | K10 | 1.30E-07 | N2840 |
| MAPK11 | N | NV1651 | K4 | 1.30E-07 | N2540 |
| ABL1 | N | NV1011 | K4 | 1.30E-07 | N2540 |
| BRAF | C | NV2481 | K10 | 1.00E-06 | N2840 |
| CAMK1 | N | NV2531 | K9 | 6.60E-07 | N2830 |
| JAK1 | C | Kind gift of Promega | K10 | 2.50E-07 | N2840 |
| KIT | C | NV1491 | K4 | 6.30E-08 | N2540 |
| LRRK2 | C | NV3401 | K9 | 8.30E-09 | N2830 |
| PRKCQ | C | Kind gift of Promega | K10 | 5.00E-07 | N2840 |
